# Decoding kinematic information from beta-band motor rhythms of speech motor cortex: A methodological/analytic approach using concurrent speech movement tracking and magnetoencephalography

**DOI:** 10.1101/2023.06.12.544529

**Authors:** Ioanna Anastasopoulou, Douglas O. Cheyne, Pascal van Lieshout, Blake W Johnson

## Abstract

Articulography and functional neuroimaging are two major tools for studying the neurobiology of speech production. Until now, however, it has generally not been feasible to use both in the same experimental setup because of technical incompatibilities between the two methodologies. Here we describe results from a novel articulography system dubbed Magneto-articulography for the Assessment of Speech Kinematics (MASK; Alves et al., 2016), which is technically compatible with magnetoencephalography (MEG) brain scanning systems. In the present paper we describe our methodological and analytic approach for extracting brain motor activities related to key kinematic and coordination event parameters derived from time-registered MASK tracking measurements (Anastasopoulou et al., 2022). Data were collected from ten healthy adults with tracking coils on the tongue, lips, and jaw. Analyses targeted the gestural landmarks of reiterated utterances /ipa/ and /api/, produced at normal and faster rates (Anastasopoulou et al., 2022; Van Lieshout, 2007). The results show that (1) Speech sensorimotor cortex can be reliably located in peri-rolandic regions of the left hemisphere; (2) mu (8-12 Hz) and beta band (13-30 Hz) neuromotor oscillations are present in the speech signals and contain information structures that are independent of those present in higher-frequency bands; and (3) hypotheses concerning the information content of speech motor rhythms can be systematically evaluated with multivariate pattern analytic techniques. These results show that MASK provides the capability, for deriving subject-specific articulatory parameters, based on well-established and robust motor control parameters, in the same experimental setup as the MEG brain recordings and in temporal and spatial co-register with the brain data. The analytic approach described here provides new capabilities for testing hypotheses concerning the types of kinematic information that are encoded and processed within specific components of the speech neuromotor system.

## Introduction

In recent years, systematic studies of speech motor control in the human brain have significantly expanded our understanding of the neural foundations of expressive speech. Converging evidence now points to a comprehensive re-evaluation of conventional and long-held theoretical models of speech production (see recent reviews by Hickok & Venezia, 2023 and Silva et al., 2022). This re-evaluation is ongoing and rapidly evolving, but there is an emerging shift away from the traditional Wernicke-Geschwind model’s emphasis on Broca’s region in the left hemisphere, and towards a greater recognition of the computational roles and network connections of various premotor, motor, sensory, and insular regions of the cerebral neocortex (Hickok & Venezia, 2023).

Much of the information that informs these new models comes from non-invasive neuroimaging techniques, predominantly functional magnetic resonance imaging (fMRI) (Bohland & Guenther, 2006; Peeva et al., 2010; Pang et al., 2011; Behroozmand et al., 2015; Rong et al., 2018; Tourville et al., 2019; Heim & Specht, 2019) and associated techniques including diffusion tensor imaging (DTI) (Catani & Forkel, 2019; Chang et al., 2020; Janssen et al., 2022).

Non-invasive electrophysiology methodologies using electroencephalography (EEG) and magnetoencephalography (MEG) have added important detail regarding the timing of neuronal processing events (Munding et al., 2016; Salmelin et al., 2019; Leckey & Federmeier, 2019). Finally, recent years have provided an increasing amount of very highly detailed electrophysiological evidence from invasive electrophysiological (electrocorticography; ECoG) recordings of speech motor regions in neurosurgical patients (Bouchard et al., 2013; Ramsey et al., 2018; Chartier et al., 2018; Silva et al., 2022).

Current evidence shows that spoken language processing draws on a complex set of neural computations performed in a widely distributed set of brain regions (Levelt et al., 1998; Munding et al., 2016; Carota et al., 2022). These computations range from abstract aspects of semantics and syntactics to the low-level sensorimotor processes that directly control and modulate the overt movements of speech articulators of the peripheral vocal tract (Indefrey & Levelt, 2000; Tong et al., 2022). Experimental and clinical protocols for mapping of expressive speech centres employ a wide variety of speech tasks according to their specific experimental or clinical aims (Salmelin et al., 2019). Speech tasks can be variously deployed to emphasise different aspects of spoken language processing: Story listening, object naming, rhyming, and covert word production invoke relatively more abstract linguistic processes and have been shown to reliably activate distributed areas of prefrontal, temporal and parietal cortex, including Broca’s area in the left hemisphere (Bowyer et al., 2005; Doesburg et al., 2016; Kadis et al., 2011; Youssofzadeh & Babajani-Feremi, 2019; Correia et al., 2020); while in contrast, non-word/pseudoword tasks are intended to limit the requirements for semantic, syntactic and attentional processing and elicit neural activity that is more restricted to brain regions associated with sensorimotor processes (Frankford et al., 2021; Kearnery & Guenther, 2019).

The subject of the current paper is set within the context of speech motor control: the phonological, phonetic, and sensorimotor processes that control and/or modulate the neuromuscular output to the articulators. In this context, an important methodological limitation of current neuroimaging research is that, with rare exceptions (Chartier et al., 2018; Mugler et al., 2018; Ouyang et al., 2016), researchers obtain little or no information about the actual movements of said articulators. This is a fundamental limitation in light of evidence that speech (and other movements) is encoded in the form of kinematic movement trajectories in neurons in primary motor cortical neurons (Chartier et al., 2018; Conant et al., 2018; Kolasinski et al., 2020). Such essential information is technically difficult to obtain for crucially important articulators (such as the tongue) which are located out of the line of sight within the oral cavity. Unfortunately, specialised electromagnetic and ultrasound articulography techniques that are capable of non-line-of-sight speech tracking are technically incompatible with the scanner environments used for functional imaging with fMRI and MEG (Anastasopoulou et al., 2022).

In the following, we describe our method for linking speech kinematics to brain activity using a novel MEG setup. This setup enables us to simultaneously and accurately measure speech movements and brain function. The system, termed Magnetoencephalography for Assessment of Speech Kinematics (MASK, Alves et al., 2016), can track the independent motion of up to 12 lightweight coils that are similar in size and shape to the tracking coils used in conventional electromagnetic articulography (EMA). In contrast to the passive induction coils used in EMA, MASK coils are actively energized by sinusoidal currents, and their associated magnetic fields are measured by the MEG sensors.

In a previous paper we have described in detail movement parameters (amplitude, duration, velocity) and interarticulator phase relationships derived from direct MASK measurements of articulator movements (Anastasopoulou et al., 2022). We have demonstrated that MASK reliably characterizes key kinematic and movement coordination parameters of speech motor control, achieving a resolution comparable to standard electromagnetic articulography devices. In the present work, we proceed to describe our methodology for establishing a mapping between MASK-derived kinematic parameters and MEG-derived brain activities.

## Methods

### Participants

Ten healthy adults participated in this study (4F; mean age 32.5, range 19.7-61.8; all right-handed as assessed by the Edinburgh Handedness Inventory). All participants were fluent speakers of English; Nine were native English speakers, one participant’s first language was Mandarin. All procedures were approved by the Macquarie University Human Research Ethics Committee.

### MEG scans

Speech tracking data and neuromagnetic brain activity were recorded concurrently with a KIT-Macquarie MEG160 (Model PQ1160R-N2, KIT, Kanazawa, Japan) whole-head MEG system consisting of 160 first-order axial gradiometers with a 50-mm baseline (Kado et al., 1999; Uehara et al., 2003). MEG data were acquired with analogue filter settings as 0.03 Hz high-pass, 1000 Hz low-pass, 1000 Hz sampling rate and 16-bit quantization precision in a magnetically shielded room (Fujihara Co. Ltd., Tokyo, Japan). Measurements were acquired with participants in a supine position (as the KIT MEG dewar configuration is fixed in the supine position). This positioning is in contrast to conventional standalone articulography setups which are used with participants in an upright seated position. We note that recent research findings have indicated that there is no significant distinction in tongue pressure between participants in a supine position and those in an upright position (Dietsch & Cirstea, 2013).

### Structural scans

T1-weighted anatomical magnetic resonance images (MRIs) were acquired for all participants in a separate scanning session using a 3T Siemens Magnetom Verio scanner with a 12-channel head coil. Those anatomical images were obtained using 3D GR\IR scanning sequence with the following acquisition parameters: repetition time, 2000 ms; echo time, 3.94 ms; flip angle, 9 degrees; slice thickness, 0.93 mm; field of view, 240 mm; image dimensions, 512 × 512 × 208.

### Procedure

Five head position indicator coils (HPI) were attached on the head in an elastic cap, and their positions were measured at the beginning and at the end of the experiment, with a maximum displacement criterion of < 5 mm in any direction. The coils’ positions with respect to the three anatomical landmarks (nasion, right and left preauricular landmarks) were measured using a handheld digitiser (Polhemus FastTrack; Colchester, VT).

MASK coils were placed at mid-sagittal positions on the vermilion border of the upper lip (UL) and lower lip (LL), the tongue body (TB; 2 cm from the tongue tip) and the lower incisor (JAW) sensor which was attached to a thin thermoplastic mould (Van Lieshout et al., 2007). Tongue sensors were attached with surgical glue (Epiglu, MajaK Medical Brisbane; Australia), while lip sensors were attached with surgical tape. Tracking coils were driven at frequencies higher than 200 Hz, which allowing separate coil fields from brain activities that primarily occur at frequencies lower than 100 Hz. Line-of-sight is not required for MASK measurement, permitting tracking of all oral articulators, including the tongue. MASK evaluates coil positions every 33 ms for movement tracking at rates up to 50 cm/s (Alves et al., 2016). Spatial accuracy depends on the distance of the tracking coils from the MEG sensor array. For coils that are close to the array (like those on the tongue), the accuracy is less than 1 mm relative position error, similar to the standard MEG head position indicator coils. However, for coils that are more distant from the helmet sensor array (like those on the lower lip), spatial accuracy decreases non-linearly to approximately 1-2 mm.

An experimental control computer with scripts programmed in Presentation® software (Version 23.0, Neurobehavioral Systems, Inc., www.neurobs.com) was used to control display of experimental instructions which were projected onto a display screen placed 100 cm above the head of participants.

160 channel MEG data were simultaneously relayed to the MEG data acquisition computer and to a separate MASK processing computer which calculated coil positions from the MEG signals with an offline localisation algorithm, using a computational approach similar to those used in conventional MEG for localising fiducial coils with respect to MEG sensors (Wilson, 2004), with modifications to optimise performance of the smaller MASK tracking coils (Alves et al., 2016; More, 1978).

Acoustic speech signals were recorded with a directional microphone placed on the wall of the magnetically-shielded room and were digitised in an auxiliary channel of the MEG electronics with the same sample rate (1000 Hz) as the MEG recordings, and was relayed to both the MEG acquisition computers and the MASK processing computer. The MASK-processed speech tracking signals were subsequently combined with the MEG datasets stored on the acquisition computer. Due to differences in the internal clocks of the MEG and MASK computers, it was necessary to time-align the two datasets by applying the MATLAB *alignsignal* script to the acoustic channels of the MEG and MASK datasets.

An additional high sample rate speech recording was obtained with an optical microphone (Optoacoustics, Or-Yehuda, Israel) fixed on the MEG dewar at a distance of 20 cm away from the mouth of the speaker; and digitised on the experimental control computer using a Creative sound blaster X-Fi Titanium HD sound card (Creative, Singapore) with 48 kHz sample rate and 24-bit quantization precision. The 48 kHz acoustic recording was aligned and integrated with the MEG and MASK datasets as described above.

### Experimental protocol

Participants performed four speech production tasks and one manual button press task (see Figure 1). Speech productions were non-word disyllabic sequences with a V1CV2 structure, /ipa/ and /api/, and each was produced in a reiteration fashion at normal and faster rates. The tongue and lip gestures for /ipa/ and /api/ are reversed in phase, providing a robust behavioural contrast in terms of interarticulator coordination (Anastasopoulou et al., 2022; Van Lieshout et al., 2007). Variations in speech rate were used as a control variable to examine the intrinsic stability of the coordination (Kelso 1986, Van Lieshout et al., 1996). Asking participants to change their speaking rate (frequency of executed movements) is a typical characteristic of studies which investigate coordination dynamics (Kelso, 1995, Van Lieshout et al., 1996). The same reiterated stimuli have been used in previous studies investigating speech motor control strategies in normal and in disordered populations (Van Lieshout et al. 1996; Van Lieshout et al. 2002; Van Lieshout et al. 2007; Van Lieshout, 2017). Nonword stimuli with no linguistic information avoid familiarity issues (Van Lieshout, 2017) and have been widely used in the literature to investigate normal and pathological function in speech motor control (Murray et al., 2015; Case & Grigos, 2020).

**Figure 1.**
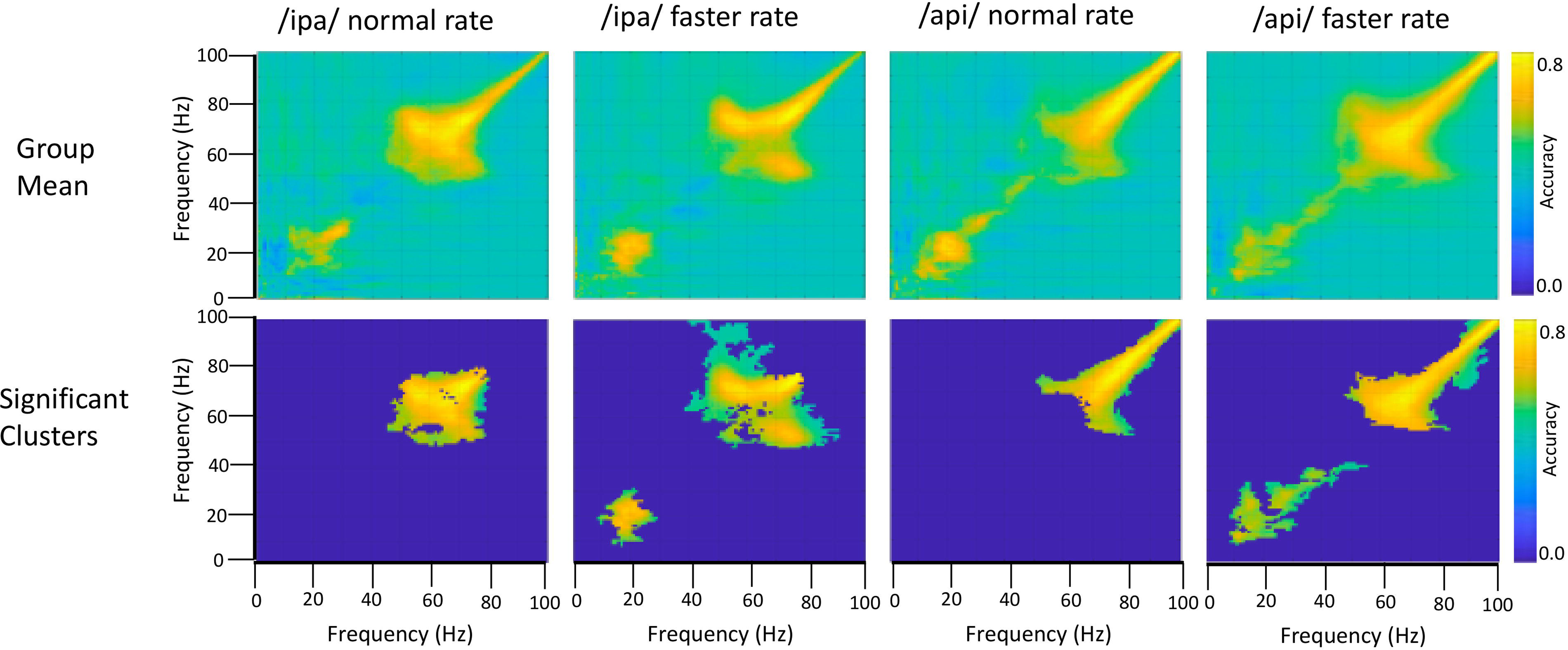
Experimental procedures. A. Speech task. Instructions were displayed for 30s, followed by an intertrial interval lasted for 15s and 5s fixation cross ‘+’ and breath intake in preparation for the speech production trial set. During a trial set participants produced the indicated nonword in a reiterated fashion for 12s. 10 consecutive trial sets were performed for each nonword stimulus. B. Button press task. Instructions were displayed for about 30s followed by a fixation cross, during which participants performed self-paced button pressed with the index finger of their dominant (right) hand at a rate of about 1 per 2 seconds for a total of about 90 trials.

The experimental procedure for the speech task is illustrated in Figure 1A. Participants were presented with a fixation cross (1 sec duration) on a display screen duration which they were instructed to take a deep breath. The fixation cross was replaced with a stimulus nonword for 12 sec. Participants were required to continuously utter productions during the exhalation of the single breath intake until the stimulus nonword disappeared from the screen. For the normal rate, participants were instructed to produce utterances at a comfortable rate as they would do while conversing with a friend. For the faster rate, they were instructed to produce utterances as fast as possible while maintaining accuracy (Van Lieshout et al. 2002). Following Van Lieshout (2007), we refer to the reiterated productions generated within the span of a single exhalation as a “trial set”. A respite of 4 seconds, referred to as the “inter trial set interval” followed each trial set. At the end of the intertrial set interval, a fixation cross appeared for one second to cue the intake of breath for the next trial set. Participants generated about 10 individual productions in each normal rate trial set and about 12 individual productions in each faster rate trial set. Participants performed 10 trial sets of each utterance and each rate, for a total of 40 trial sets. A pre-experiment training session exposed participants to recorded productions of a female native Australian speaker. Participants repeated the modelled productions with feedback on incorrect speech productions, rate and head or eye movements. They were allowed to blink their eyes between the trial sets but were instructed to avoid these during trial sets.

Participants also performed manual button presses on a fibre optic response pad (Current Designs, Philadelphia) with the index finger of their dominant hand at a self-paced rate of about 1 per 2 seconds for 180 sec (see Figure 1B) (Cheyne et al., 2008; Cheyne et al., 2014; De Nil et al., 2021; Johnson & He, 2019).

## Analyses

Data analyses proceeded in four main phases:

1. Analyses of MASK speech movement signals to characterise speech kinematic profiles;
2. MEG source reconstruction to identify location of speech motor cortex;
3. Extraction of MEG-time frequency spectrograms from source-localised speech motor cortex, followed by and multivariate pattern analysis of speech-relevant brain rhythms.
4. Mapping of speech kinematic profiles onto source- and frequency-constrained MEG data, via representational similarity analysis (RSA).

For the purposes of clarity, we present details of each set of analytic methods along with their results, organised according to these four analytic phases.

### Analysis of MASK-derived speech movement signals

## Methods

The raw MASK position data were head motion corrected using MASK coils placed at fiducial landmarks (nose, left and right ear) and transformed from the MEG coordinate system into the occlusal plane frame of reference such that motion signals could be measured relative to a midsagittal plane defined by the x (posterior-anterior) and z (inferior-superior) axes relative to the bite plane origin. These transformed signals were then analysed using EGUANA software (Henriques & Van Lieshout, 2013; Van Lieshout, 2021) to derive signal amplitude and phase for selected articulators and speech gestures. Speech errors (e.g. substitutions, lengthy pauses) were identified from examination of the tracking and acoustic signals, and subsequent analyses were focused on accurate productions (Case & Grigos, 2020).

/ipa/ and /api/ productions involve specific movements of the lips and tongue. To create the voiceless stop /p/ sound, a bilabial closure (BC) gesture is used. The two tongue body constriction gestures (TB) are used to produce the sounds /i/ and /a/. The BC gesture was calculated using the two-dimensional (x-y) Euclidian distance of the upper and lower lip positions, while the tongue body gesture was derived from the two-dimensional (x-y) Euclidian distance of the tongue body and the nasal reference coil, as described in Van Lieshout et al. (2007). The kinematic and coordination parameters were computed using the methods described in (Anastasopoulou et al., 2022; van Lieshout et al., 2002; van Lieshout et al., 2007).

The opening and closing movements of each cycle were identified using the minimum and maximum vertical position of the gestural and articulatory signals (Van Lieshout, 2017). The amplitude levels of the opening and closing movements at 10% and 90% were determined for each individual cycle. The individual gestural and articulatory signals which were measured by the MASK (see Anastasopoulou et al., 2022) were analysed with custom MATLAB scripts to determine the times which were corresponding to the 10% and 90% amplitude levels of each opening and closing movement. Once the individual landmarks were determined, they were brought into time register with MEG data by aligning the acoustic signal recorded by the MASK acquisition setup and the acoustic signal recorded in the MEG acquisition computer, as shown in Figure 2.

**Figure 2:**
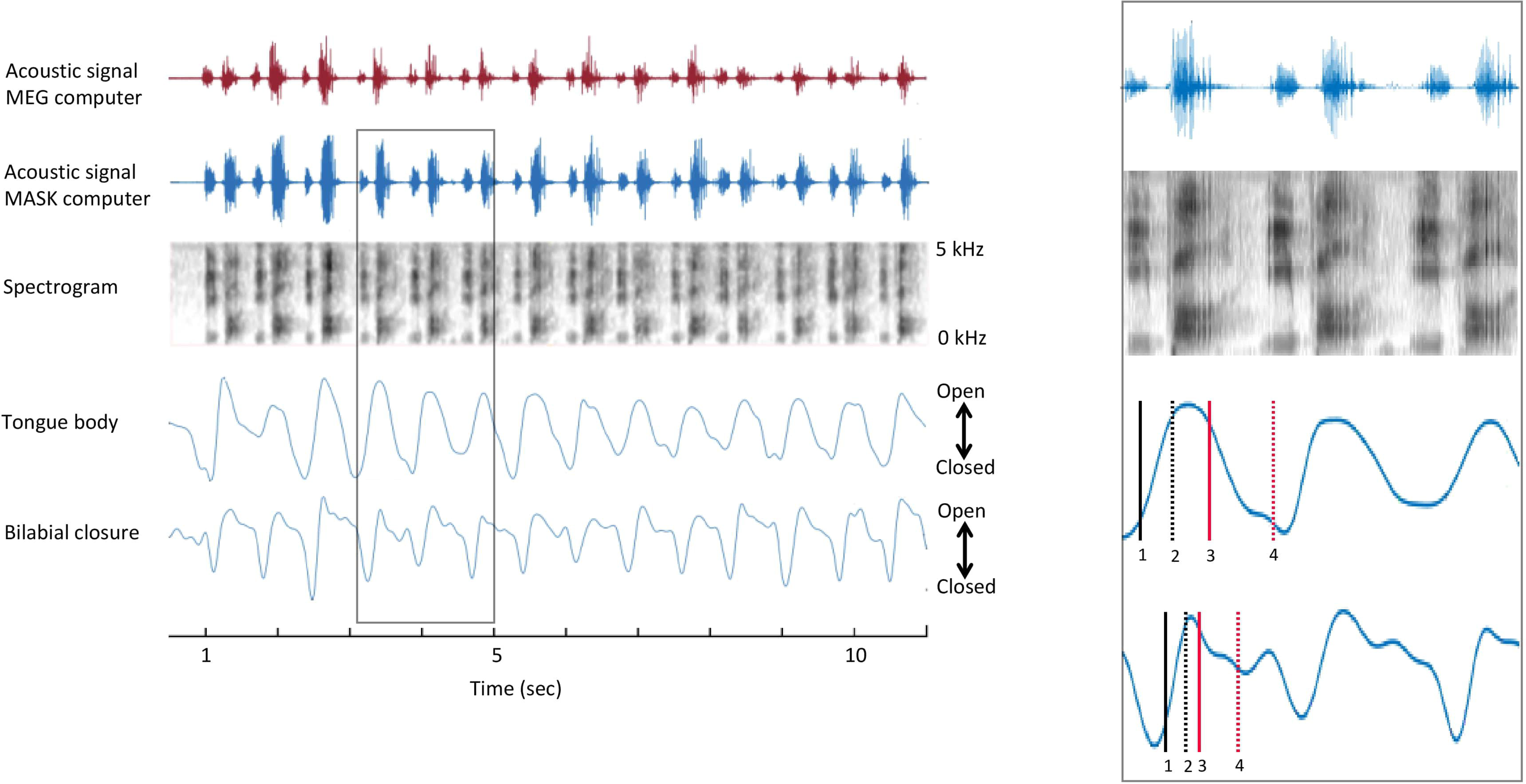
Temporal alignment of MEG and MASK signals. MASK articulatory signals are brought into register with the MEG brain data using the MATLAB *alignsignals* function on the MEG auxiliary channel acoustic channel and the high-fidelity acoustic recording (top and second rows). Inset shows enlargement of rectangle bounded area in main figure. 1 = onset of opening movement; 2 = offset of opening movement; 3 = onset of closing movement; 4 = offset of closing movement.

## Results

### 1B. MASK tracking signals

Figure S1 displays acoustic recordings and tracking signals of the tongue body (TB) and bilabial closure (BC) from Participant 1 for a single trial of each of the four speech production tasks. The participant produced 14 utterances of /ipa/ and /api/ at a normal speaking rate and 18-19 utterances at a faster rate.

The contrast between the mirrored positions of the tongue and lips in /ipa/ and /api/ is clearly observed in the MASK measurements of tongue and lip gestures. Peaks and valleys in Figure 3 indicate the high and low positions achieved by the BC and TB gestures during the production of /api/ and /ipa/. Valleys occur during the bilabial constriction gesture and the tongue body gesture for /i/, while peaks occur for the tongue body gesture of /a/. (We note that these positions are Euclidean distances relative to the nasion. Within this reference frame “low a” is a peak, and “high /i/” is a valley). In /api/, the /p/ closure occurs during the upward motion of the TB, going from the low /a/ to the high /i/ position. On the other hand, in /ipa/, the /p/ closure occurs during the downward motion of the TB, going from the high /i/ to low /a/ position. The gestural movements of /ipa/ and /api/ are mirror images, with the relative timing of the motions of TB and BC gestures reversed.

**Figure 3.**
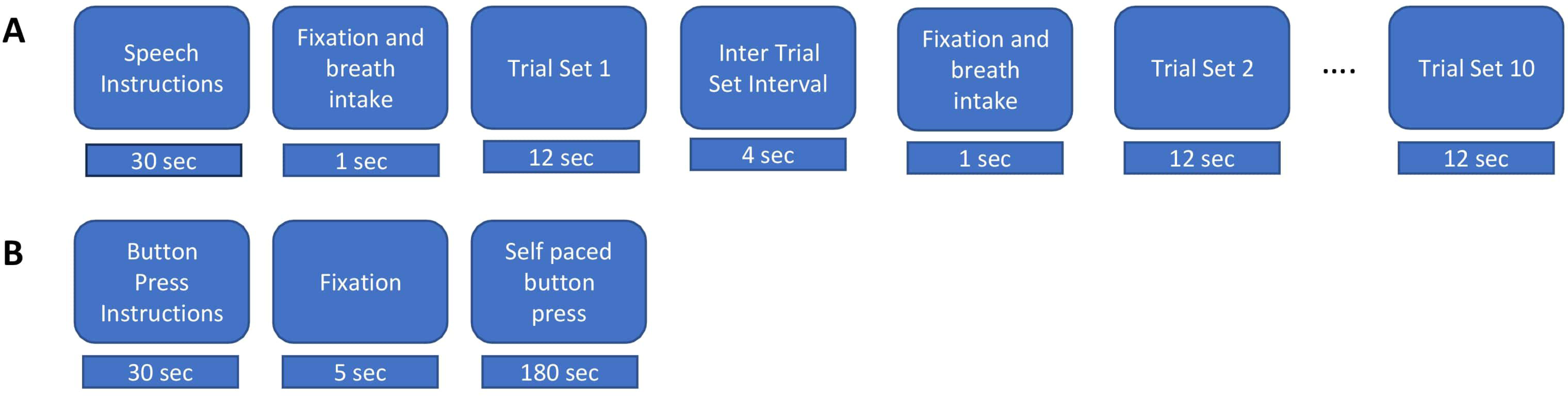
Localisation of speech motor cortex. ***Left: Anatomical Landmarks.*** 1 – Hand regions of precentral gyrus (hand knob); 2 – Hand region of postcentral gyrus; 3 – Middle precentral gyrus; 4 – Middle frontal gyrus; 5 – Rolandic fissure; 6 – precentral gyrus; 7 – postcentral gyrus. *Top right panel*: *SAM beamformer maps*. Button press task elicited activation of hand region of pre (motor) and postcentral (somatosensory) gyri. Speech task shows maximal activation in middle precentral gyrus, immediately ventral to the hand motor region of the precentral gyrus. Speech activation cluster also encompasses the middle frontal gyrus immediately adjacent to the middle precentral speech region. Bottom right panel: Time-frequency plots showing temporal evolution of oscillatory responses at virtual sensors placed at locations of cluster maxima shown above. For button press task, upward arrow indicates time of button press onset. For speech task, downward arrow shows speech trial set offset, upward arrow shows speech trial set onset.

Overall, the TB and BC tracking signals measured with MASK are entirely comparable in morphology and quality with those obtained from a conventional electromagnetic articulography setup (please see Anastasopoulou et al., 2022, for a direct comparison of MASK and EMA signals measured during the same utterances described here).

### Derived kinematic profiles

The next stage in our analysis pipeline involves generating profiles that capture the relationships between key kinematic parameters of BC and TB gestures. Specifically, we examine the amplitudes, durations, velocities, and stiffnesses of gestural movements, as these parameters are known to covary in highly consistent ways and reflect “invariant” properties of speech kinematic movements. These invariant properties are crucial in understanding the motor control of human speech (Munhall et al., 1989; Van Lieshout et al., 2007).

Figure S2 illustrates the covariation of these kinematic parameters for /ipa/ and /api/ for two participants. Our analysis shows that movement peak velocity increases as a linear function of movement amplitude, indicating that larger movement distances are associated with higher peak speeds. Furthermore, we observe comparable amplitude/velocity relationships for opening and closing movements, suggesting that these parameters are controlled similarly regardless of movement direction. This roughly linear relationship between amplitude and peak velocity is a well-established characteristic of speech kinematics, and has been described for a variety of articulators, gestures, and utterances (Ostry et al., 1983; Ostry et al., 1985). Regarding the stiffnessvs. duration relationship, our results indicate that stiffness systematically decreases as a curvilinear function of durations less than 200 ms, after which the relationship plateaus into a relatively flat line (see also Munhall et al., 1989). These findings are in line with the findings reported in Anastasopoulou et al., (2022) and consistent across the ten speakers.

### MEG source reconstruction of speech motor cortex

## Methods

Source reconstruction of brain activity was carried out using the synthetic aperture magnetometry (SAM) beamformer algorithms implemented in the BrainWave MATLAB toolbox (Jobst et al., 2018; cheynelab.utoronto.ca/brainwave). The raw KIT/Yokogawa data files were initially converted to CTF format and transformed to the CTF head coordinate system using the fiducial coil positions relative to the sensor array. Each participant’s structural MRI was then spatially coregistered with the MEG data and normalised into standard adult MNI template space using SPM12 (Wellcome Institute of Cognitive Neurology).

The onset of each speech trial set was marked using a semi-automated peakfinding function implemented in the BrainWave MATLAB toolbox which identified peak onsets in the rectified signal of the auxiliary MEG acoustic channel. In this analysis, we use an acoustic rather than speech movement index of speech trial onsets, because the aim of localising speech motor cortex is independent of the aims of our subsequent kinematic analysis, and because the acoustic onset provides ample accuracy for the SAM beamforming analysis.

Raw data were pre-filtered using a 100 Hz low pass bidirectional zero phase-shift Butterworth filter and epoched into 15 sec segments, from 10 sec prior to speech trial onset to 5 seconds after speech trial onset. Each 15 sec segment encompassed three distinct task periods: the last five seconds of the preceding trial set (-10 to – 5 sec); the inter-trial set rest period (-5 to 0 sec); and the first five seconds of the current trial set (0 to +5 sec), thereby providing maximal contrast between active (speech) and rest periods.

The current trial set and inter-trial rest period intervals were used for the SAM pseudo-T analysis window and baseline window respectively. We used a sliding active window of 1 second duration starting from 0-1000 ms (step size 200 ms, 10 steps), and a fixed baseline window of 2 seconds duration extending from -5 to -3 seconds relative to speech movement onset and a bandpass of 18– 22 Hz (centre of the beta frequency range). While beamforming analyses typically employ equal baseline and active window durations, in the present analyses we wished to use a longer baseline epoch to reduce the chance of a biased estimate of baseline power, which we reasoned is more likely to vary over time than during the active speech period (since reiterated speech is akin to a steady state). While larger order mismatches (e.g. comparing 1.0 second to 0.1 seconds) are likely to be problematic, simulation studies (Brookes et al., 2008) have shown that covariance errors are minimized, and therefore beamformer weights are stable, with at least 5 seconds of total data (a requirement that is well exceeded in the present case) and our preliminary analyses confirmed that a 2 second baseline did not affect the beamforming results relative to a 1 second baseline. The combined active and control windows were used to compute the data covariance matrix for beamformer weight calculations, while the full 15 second time window was used to compute data covariance for the virtual sensor source activity calculations.

SAM pseudo-T images were volumetrically reconstructed using a 4 mm resolution grid covering the entire brain. Group statistical analysis employed a nonparametric permutation analysis where each permutation (total = 1024) involved randomly flipping the polarity of all voxels in the group volumetric image, computing the mean image, and placing the maximum value in the image in the permutation distribution (Jobst et al., 2018; Nichols and Holmes, 2002; Chau et al., 2004; Lin et al., 2006). The value corresponding to the area under the permutation distribution for the Type I error level (alpha = .05) was used as an omnibus threshold for the entire image. The time-course of (group) source activity was then computed as the output of the beamformer with optimized orientation (“virtual sensor”) and plotted as time-frequency spectrograms (encompassing the entire 15 sec data epoch) to assess the temporal correspondence of beta activity with active and rest periods. To maximise the number of trials (and consequently, the signal to noise ratio) in this analysis we use trial sets from all four speech tasks (for a total of 40 trial sets). To examine individual datasets, statistically significant locations were unwarped from the group template to the same anatomical locations in the individuals and virtual sensors were computed in the same manner as described above for the group analysis.

For the motor cortex hand region localiser analysis, trials were prefiltered with a bandpass of 0-100 Hz and epoched with respect to the button press onset into 1.5 sec segments (-500 to +1000 ms), encompassing the established time course of beta-band desynchronisation (several hundred ms prior to and after the button press) and “rebound” synchronisation (several hundred ms starting about 500 ms after the button press) (see Cheyne, 2013; Cheyne et al., 2014; Johnson et al., 2016). Following the maximal contrast approach used for the speech analysis, the SAM pseudo-T analysis used a sliding active window of 200 ms duration starting from 600-800 ms, (step size 10 ms, 10 steps), a fixed baseline window from 0 to 200 ms, and bandpass of 18-22 Hz. The full 1.5 second epoch was used to compute the data covariance matrix for beamformer weight calculations. Volumetric reconstruction used the same grid employed for the speech analysis.

In all individuals SAM source reconstruction resulted in robust peaks centred on the hand knob of the left precentral gyrus (Yousry et al., 1997), and a smaller mirror source centred on the right hemisphere homologue. The left hemisphere virtual sensor source activity was then computed and plotted as a time-frequency spectrogram (encompassing the entire 1.5 sec data epoch) to assess the correspondence with the established time course of beta band activity associated with the manual button press task (Cheyne, 2013; Cheyne et al., 2014; Johnson et al., 2020).

## Results

Group statistical analyses showed that a single significant SAM beamformer cluster (Figure 3) in the left hemisphere, encompassing the middle portion of the prefrontal gyrus (mPFG) and the immediately adjacent region of the middle frontal gyrus (MFG), both established areas of speech motor control (Silva et al., 2022). The anatomical localisation of the mPFG is well-supported by comparison with the SAM beamformer map for the button press task, which shows a cluster maximum in the hand knob of the immediately dorsal region of precentral gyrus. Physiological activities at the locations of the SAM beamformer cluster maxima are visualised in the “virtual sensor” time frequency plots below their respective brain maps. For the button press task the time-frequency plot shows the well-established pattern of beta-band (13-30 Hz) desynchronisation, starting several hundred ms before the button press, persisting for several hundred ms after, and followed by a “rebound” beta synchronisation at about 600-700 ms after the button press.

A comparable pattern of beta band activity is evident in the speech virtual sensor plot, keeping in mind the different time scales (1.5 sec for button press, 15 sec for speech) and movement requirements (a single punctate button press versus 10 seconds of steady state, reiterated speech) of the two tasks. Beta band desynchronisation begins several hundred ms before speech onset and persists for the duration of the speech movements. Note that in this plot the baseline of the colour scaling (the five second inter trial set rest period) was chosen to emphasise event-related desynchronisation. Baselining to the speaking portions of the epoch will emphasise the event-related synchronisation during the rest period.

Taken together, the results of the localisation procedure provide a focussed and well-grounded target for subsequent analyses that can incorporate the kinematic and coordination parameters derived from MASK. The plausibility of the mPFG/MFG target is well-supported by the time-frequency characteristics of the virtual sensor and its anatomic location immediately ventral to the established landmark of the PFG hand knob, independently localised with data from the button press task: a task that has been long established to provide highly reliable beta-band activations located in the hand regions of the sensorimotor cortices (e.g. Cheyne et al., 2014).

### Extraction and pattern analysis of source-localised MEG time-frequency spectrograms

Our aims in this analysis were twofold: (1) to perform a “time-frequency classification” to determine if the trial-by-trial time-frequency data derived from the speech motor cortex virtual sensor contains information that is able to discriminate between the neural activities associated with speech and non-speaking epochs, and, if so, to determine if the discriminative information is confined to a specific frequency range^1^ (Treder, 2020); and (2) to perform a “frequency generalization/cross decoding” analysis to assess whether speech related neural responses may be contaminated with by non-neural artefacts associated with overt speech movements.

## Methods

For each production task, continuously recorded MEG signals were pre-filtered with a bandpass of 0-100 Hz and 50 Hz notch filter and segmented into 4 second epochs (-2 sec to +2 sec) using the onset of the BC opening movement for each speech task as time zero. For each speech condition and participant from 75 to 250 BC opening movements were identified within the 10 trial sets (average 175).

We note that in this analysis only activity around the single BC movement at time zero is precisely time-locked and activations around other movements will necessarily be blurred due to intra- and inter-individual variations in speech rates. The present analyses therefore simply rely on the presence of task-related modulation of activity during the speech condition but not in the rest condition. This consideration also does not affect inferences from the frequency generalisation analysis since task-related neural modulations and artefacts due to overt movements are affected identically.

Data epochs were subsequently truncated to 3 seconds (-1.5 sec to + 1.5 sec) to remove edge effects from the frequency analysis. Using the speech motor cortex coordinates derived from the speech localiser for a (single) virtual sensor, time-frequency spectrograms were generated for each individual trial and the resulting three-dimensional (time x frequency x trial) matrix was exported for classification analysis using the MVPA-Light MATLAB toolbox for classification and regression of multidimensional data (Treder, 2020).

Equivalent duration non-speaking “resting” condition epochs were derived by randomly selecting epoch-reference time-points from the inter-trial set periods of the MEG data. For each speaking condition and participant, an equal number of resting condition trials was epoched.

We performed a searchlight analysis using a binary linear discriminant analysis (LDA) classifier and a metric of “accuracy” (fraction of correctly predicted class labels, range = 0 – 1), with training parameters of five folds and five repetitions. Group level statistics were performed using nonparametric permutation testing and cluster corrections for multiple comparisons (Maris & Oostenveld, 2007). As implemented in the MVPA-Light toolbox (Treder, 2020), group statistics are calculated by testing obtained accuracy values against the null value of 0.5. To create a null distribution, data is permuted by randomly swapping the obtained value and its null value.

## Results

Figure 4 shows source-localised time-frequency spectrograms for individual participant S1119. The speech-condition spectrograms show clear speech rate related modulation of circa 20 Hz beta-band activity in all speech conditions. A clear and distinct pattern of circa 10 Hz mu band activity is also observable for both of the /api/ productions. Both the beta and mu-band rhythms are well-known and established rhythms of the central motor cortices (Cheyne, 2013; Cheyne et al., 2014; Pfurtscheller et al., 2003).

**Figure 4.**
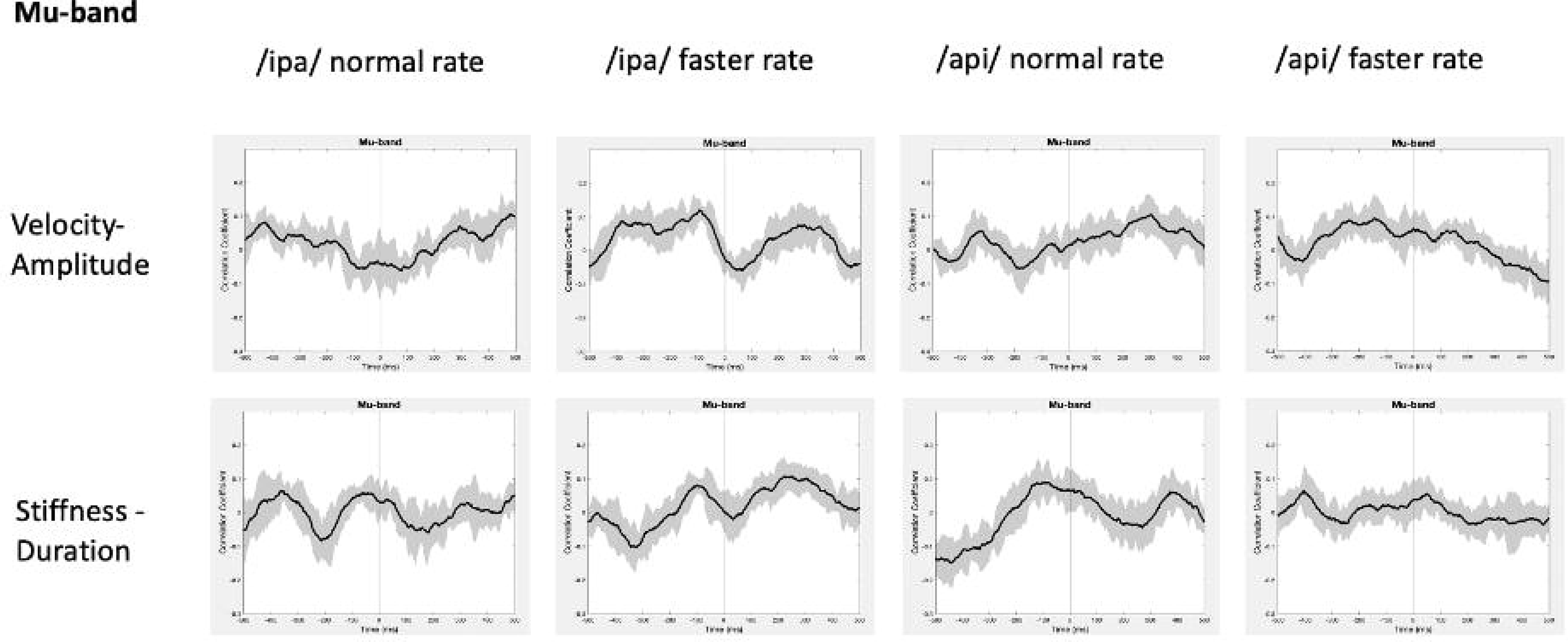
Time-frequency characteristics of speech and resting conditions for an individual participant. All plots show three seconds of MEG data derived from the medial frontal gyrus voxel. *Top row*. Time-frequency spectrograms during speech. Data are epoched relative to the onset of the bilabial closure opening movement. Speech rate modulated beta-band (circa 20 Hz) activity is evident in all plots, and mu-band activity is evident in several, especially the /api/ normal rate condition. *Middle row*. Spectrograms derived from the inter-trial set rest periods. Relatively continuous beta-band ERS is evident in all plots, as well as mu-band ERS in the /api/ conditions. *Bottom row*. MVPA classification results for speech versus resting conditions. High classification accuracy is quite tightly constrained to circa 20 Hz beta band, and a well-defined mu-frequency band is evident in the /ipa/ faster rate and /api/ normal rate conditions.

Substantial movement-related broadband noise is also evident in the supra-beta frequencies for all speaking conditions and, in this case, is especially prominent in the /api/ faster rate condition.

In contrast to the speaking conditions, for the non-speech resting conditions mu and beta activities are manifest as relatively continuous (unmodulated) bands of activity throughout most of the epoch, and broadband movement-related noise patterns are absent from the resting spectrograms.

The MVPA classification results of Figure 4 shows that the classification appears for both the mu and beta band rhythms, with a well-defined frequency boundary between the two rhythms that is clear and prominent in the cases of the /ipa/ faster rate and /api/ normal rate date. We note that the classifier is also sensitive to speech-movement related noise (present in the overt speech condition), particularly in the /api/ faster rate condition. Speech movement-related noise is evident as high-frequency broad-band patterns extending to circa 50-60 Hz. The broadband noise in the classification patterns is well separated in frequency from the beta/mu classifier signals and confined to frequencies above 50 Hz.

Level 2 group analysis of the speaking/resting classifier results are shown in Figure 5. The group results are entirely consistent with the individual results described above and provide clear statistical support for high classification accuracy for the mu and beta motor rhythms. The >50 Hz frequency region also shows high classification accuracy for broadband bursts associated with overt speech movements. The mu/beta frequencies are well separated by a region of low-classifier accuracy for circa 30-50 Hz frequencies, suggesting at least a lack of continuity between these frequency regions, and possibly that the underlying informational structures of the lower-frequency motor rhythms and the broadband artifact apparent at higher frequencies are functionally independent. We consider this issue more formally in the frequency generalisation analyses below.

**Figure 5.**
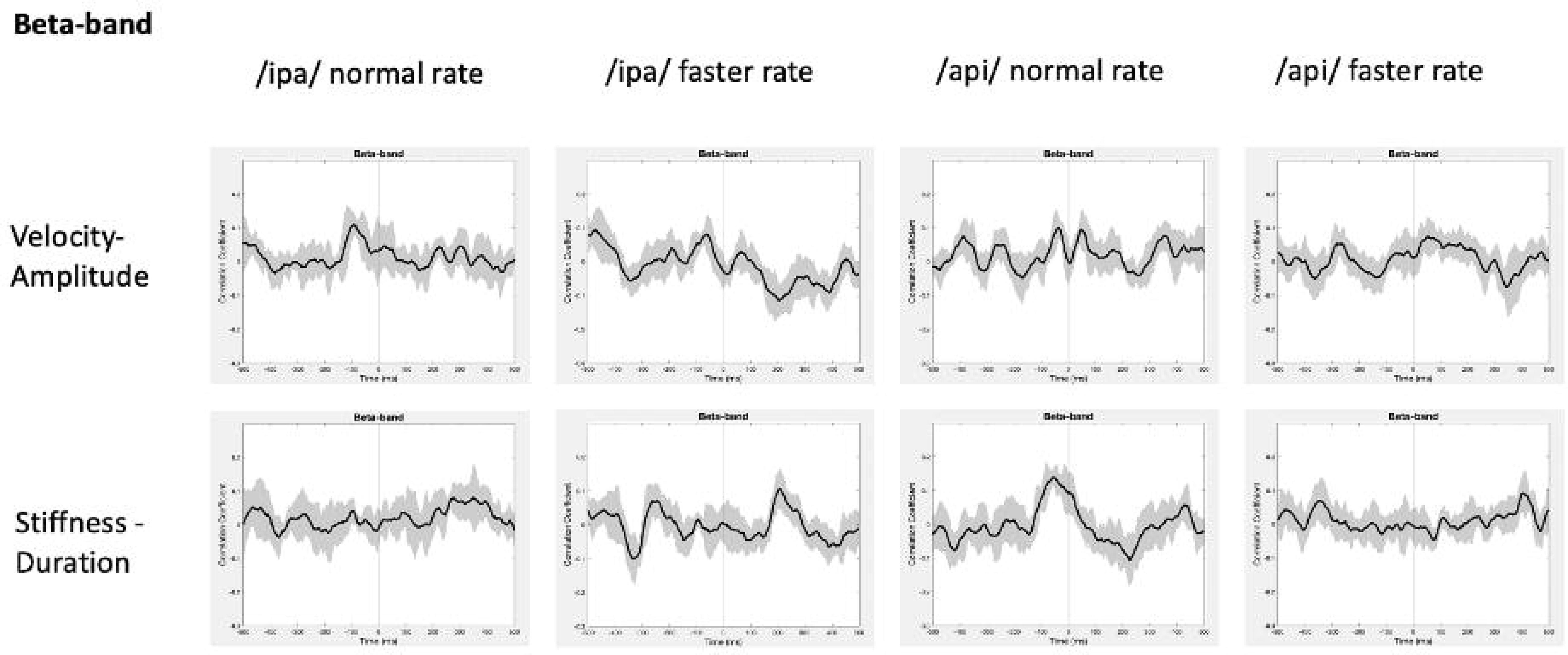
Group analysis of classifier performance for speech versus resting conditions. All plots show three seconds of MEG data derived from the medial frontal gyrus voxel. As seen in Figure 7, classifier performance is high for both speech-related high-frequency noise and beta/mu signals.

### Frequency generalisation (cross-frequency decoding)

In addition to providing an estimate of decodability for the task contrasts described above, time/frequency series decoding can be applied to provide a picture of the continuity (or discontinuity) of decoding estimates over time or frequency. This provides an important inferential advantage for further interpretation of the timing or frequency specificity of experimental effects. This “cross-decoding approach” involves training the classifier on a given time or frequency and then testing classifier performance on different times or frequencies. The logic of this approach relies on the classifier’s ability to partition multidimensional space as a basis for discriminating between experimental conditions: hence, where a classifier trained on a given time or frequency can successfully discriminate experimental classes based on other time or frequency points, one can infer that the structure of the underlying multidimensional space is similar for those two points. Conversely, in the case where cross-point decoding is unsuccessful, one can infer that the underlying multidimensional patterns are sufficiently different that the distinction between class labels determined at one point are not meaningful for discrimination at the second point (Grootswagers et al., 2017; Treder, 2020).

In the present context cross-frequency decoding enables us to more precisely address questions about possible relations between the frequency bands identified by the basic speech/rest classificational analyses described in Figures 7 and 8: (1) Do the beta/mu band signals rely on the same classification information as the high frequency signals, which are visibly contaminated with movement-related broadband signals? In this case redundancy would suggest the beta/mu signals simply contain some level of speech movement noise that is the basis for class discrimination. On the other hand, a lack of cross frequency generalisation supports the conclusion that classification relies on independent structures of information. (2) In a similar fashion, it is of interest to assess the cross-frequency generalisation between the mu and beta bands, two motor rhythms that have been frequently observed to co-occur in electrophysiological studies and can be assumed to have some functional inter-relationship (Cheyne et. al., 2014).

**Figure 6.**
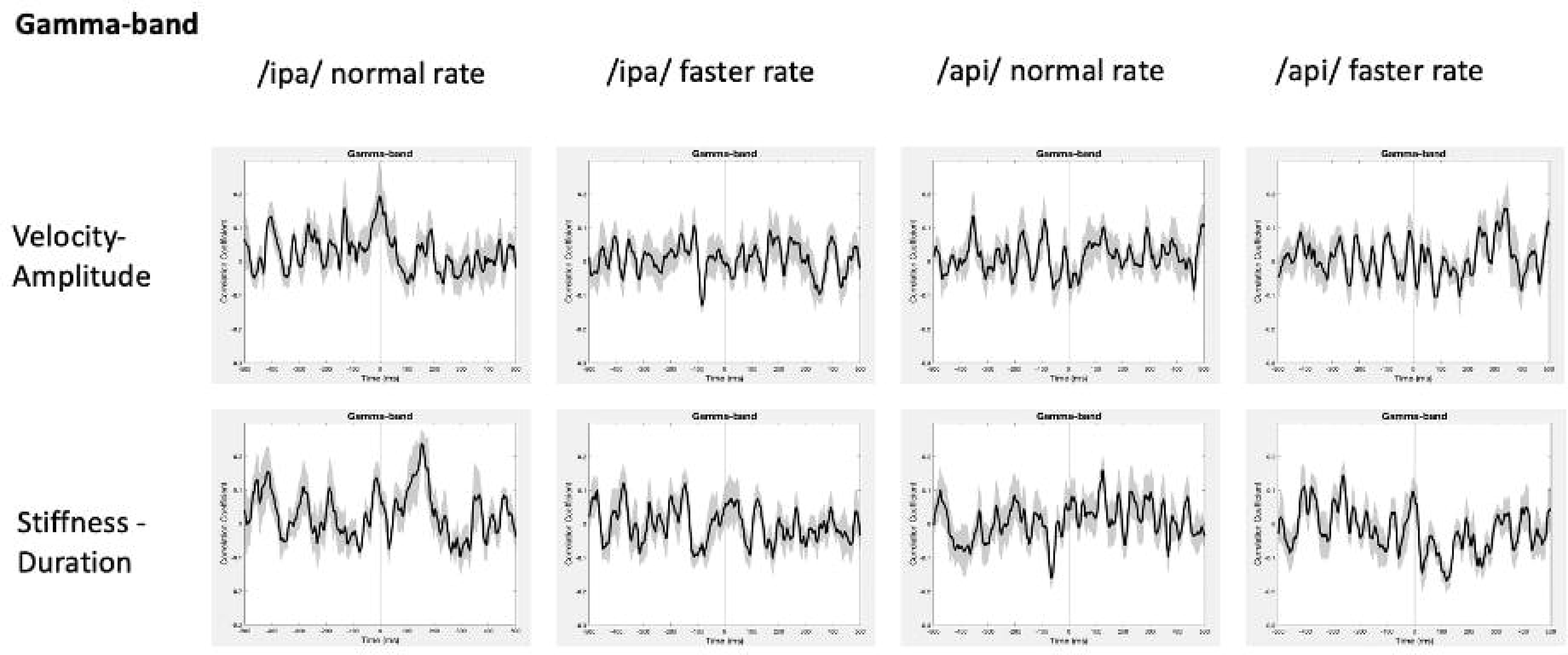
Frequency generalisation. In this analysis the classifier is trained on a given frequency and decoding performance is tested on a different frequency. This is repeated for all possible frequency pairs. The classifier results show that beta frequencies generalise to each other and to some extent to mu frequencies (bottom row, /ipa/ faster and /api/ faster). Importantly, beta/mu frequencies do not generalise to the higher frequency noise band, and conversely the noise band does not generalise to the beta/mu frequencies. Permutation-based significance tests used 500 permutations, Wilcoxin signed rank test (alpha < .05), controlled for multiple comparisons using FDR.

**Figure 7A.**
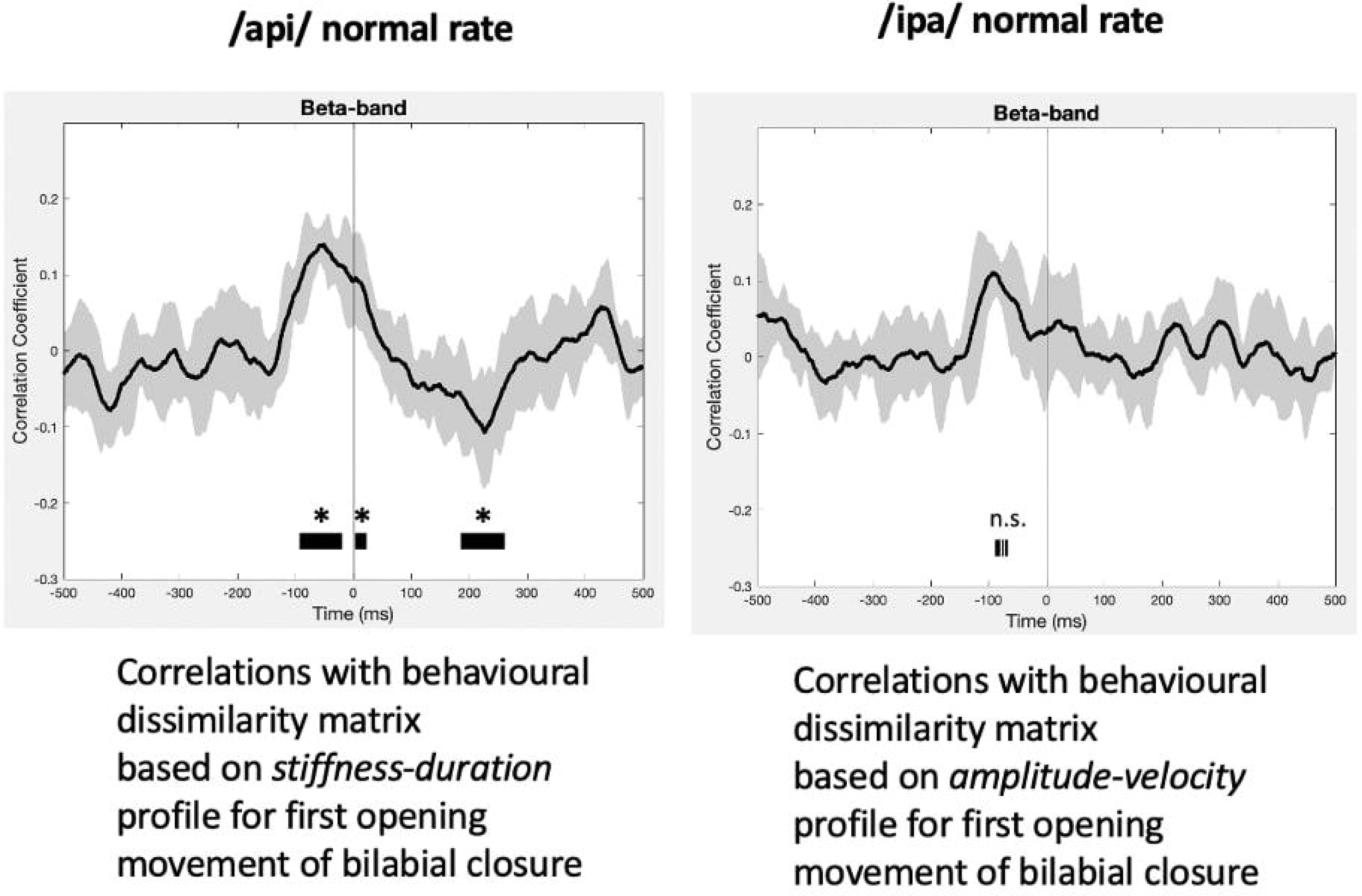
Procedure for calculation of behavioural and neural correlations. Velocity versus amplitude and stiffness versus duration profiles are partitioned into 10 blocks containing equal numbers of trials. Distances between each partition based on the partition mean coordinates are used to generate the behavioural RDMs. For each partition, the same trials of MEG data are input to MVPA classification analysis. The resulting classification (accuracy) metric is used to generate a neural RDM, for each time-frequency point.

**Figure 7B.**
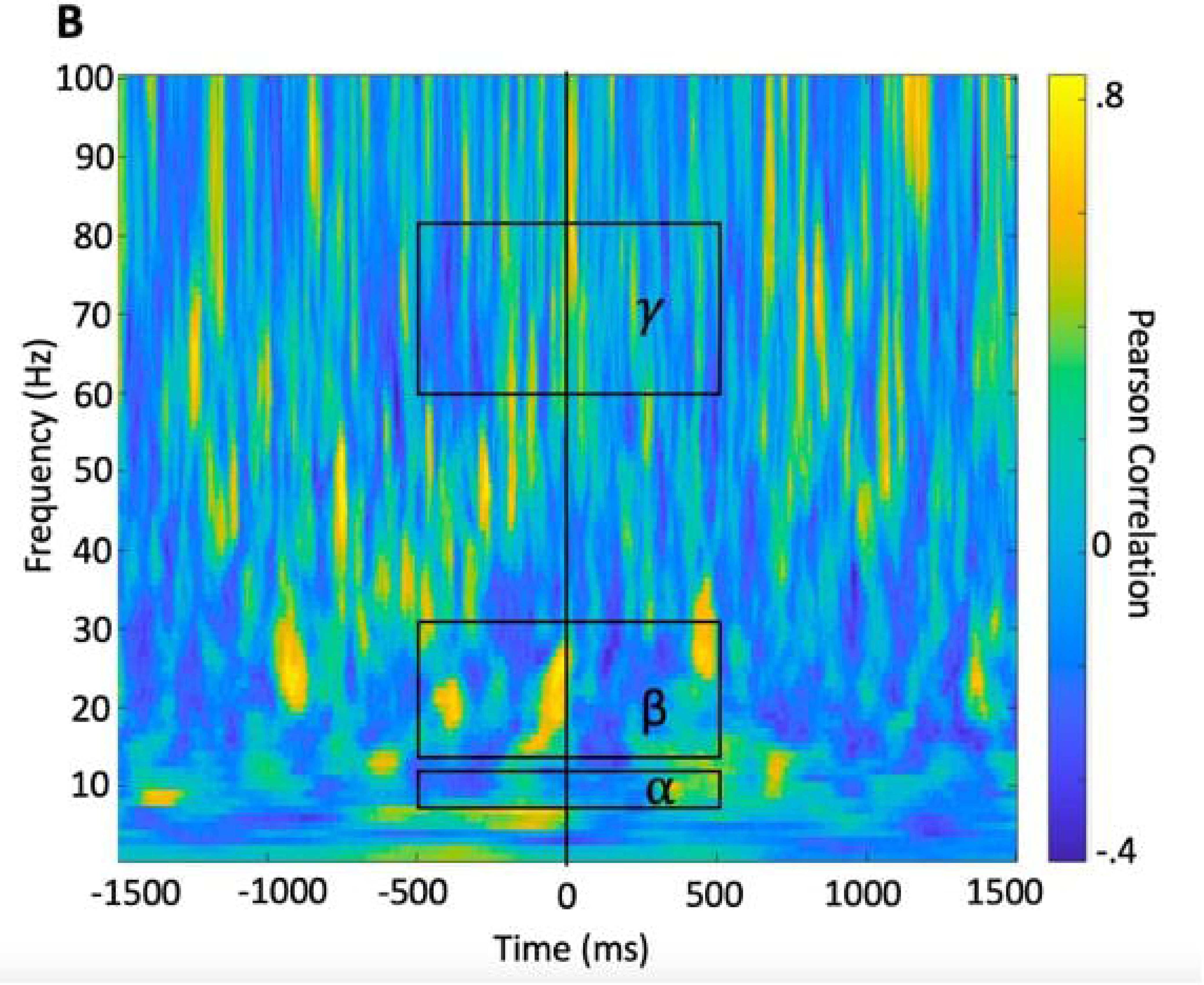
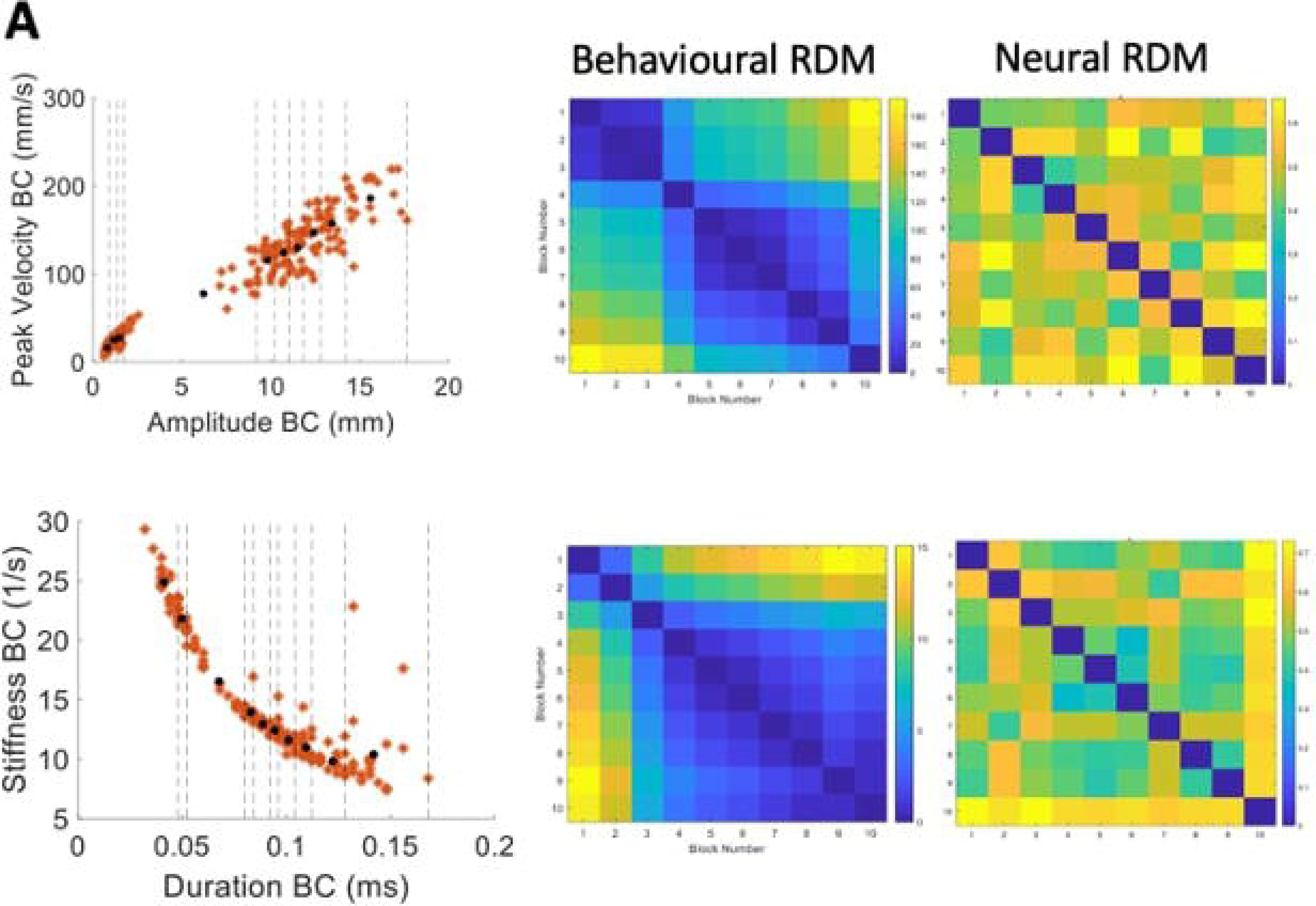
A time-frequency correlation matrix resulting from correlating the (in this case, stiffness-duration) behavioural RDM with the neural RDM for each time-frequency point. Group statistics/model evaluation are performed for the mean alpha, beta, and gamma frequency bands within the time range of -500 to +500 ms from onset of the BC opening movement.

**Figure 8.**
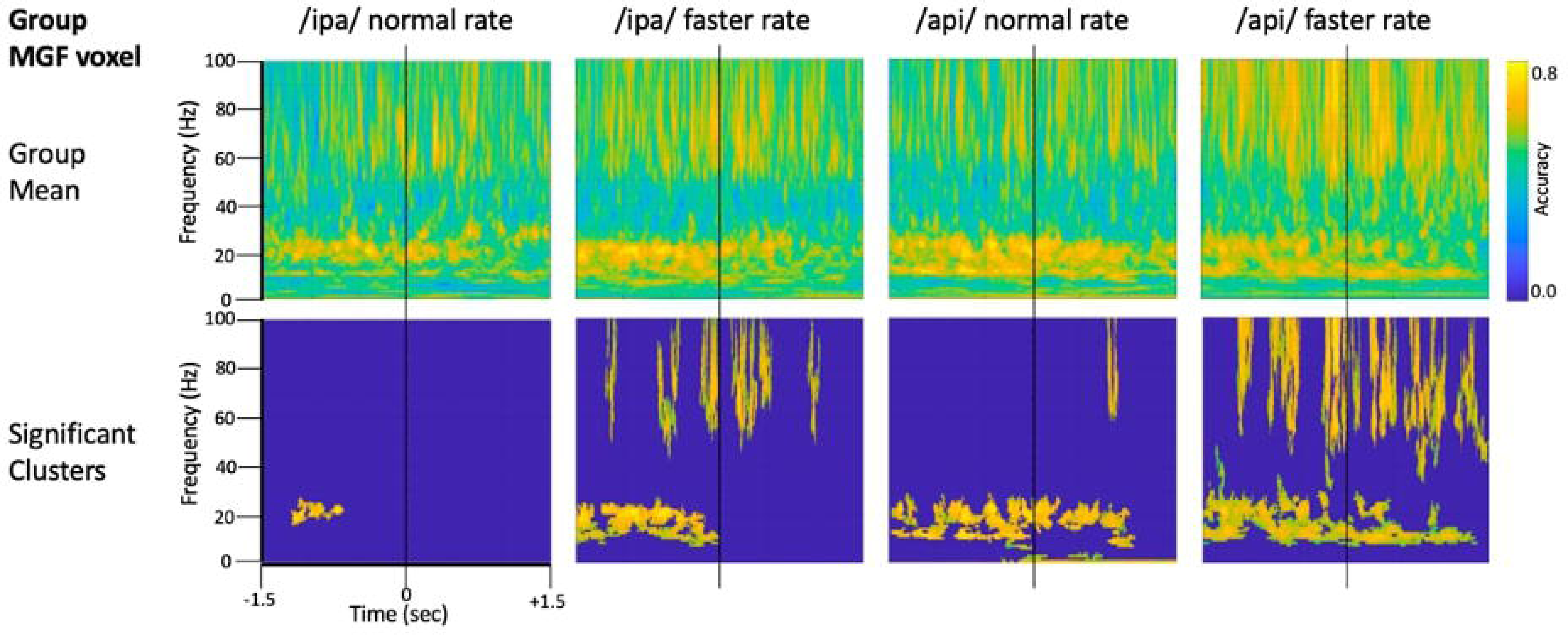
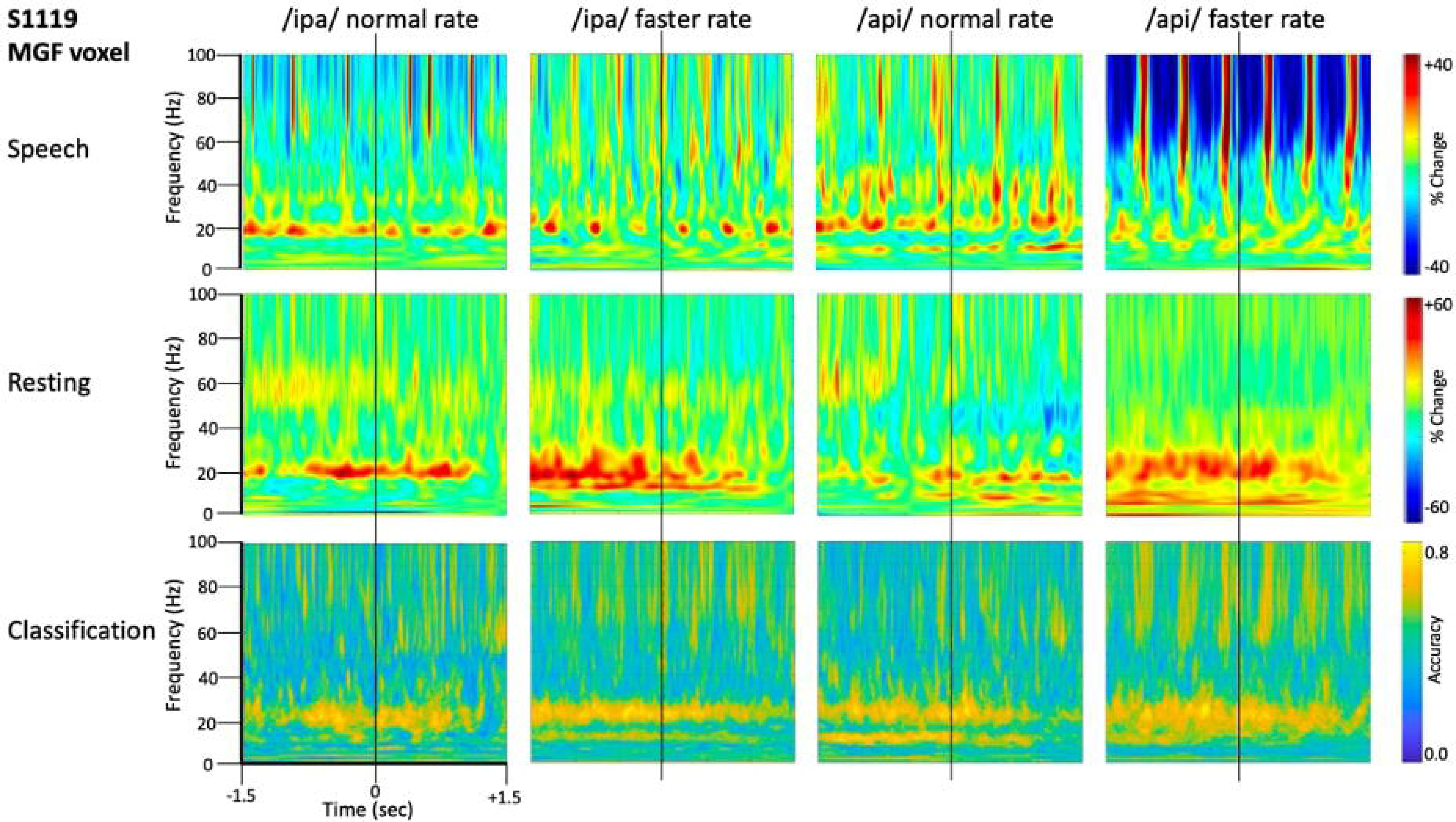
RSA model evaluation. Black lines show group mean correlations between behavioural and neural RDMs; shading shows standard errors. *Left panel*: The beta-band profile for /api/ normal rate shows three significant correlational clusters against the stiffness-duration RDM. The first positive cluster begins about 90 ms prior to onset of the first opening movement of BC. A second positive cluster occurs beginning at time zero, and a third cluster of weak *negative* correlations begins about 180 ms post movement-onset. *Right panel*: Beta correlation against amplitude-velocity time profile shows a similar positive peak circa -90 ms, although the t-value clusters do not survive cluster correction for FDR rate.

The frequency generalisation results of Figure 6 show no evidence for frequency generalisation between mu/beta, and the high frequency region which is visibly dominated by noise. To the contrary, frequency generalisation (observable as off-diagonal clustering) occurs within the sub-30 Hz mu/beta frequencies, and within the supra-60 Hz frequencies (particularly within the range of about 60-80 Hz); but the intermediate zone between mu/beta and high frequency noise (circa 30-60 Hz) exhibits a fairly strictly diagonal trajectory (for example, see group means for /api/ normal and faster rates). Classification of speech and resting conditions can clearly rely on either broadband signals that are (visibly) associated with overt speech movements, or on mu/beta modulations which show no such association: both frequency regions are prominent in the classification plots of Figures 4-5. However, the cross-frequency coding results provide clear support for the conclusion that the high frequency and mu/beta bands are discontinuous and rely on distinct patterns of multidimensional structure within their data to achieve discrimination between speech and resting data conditions.

On the second question, the group results show clear frequency generalisation between beta frequencies at circa 15-Hz and also suggest a possibly weaker generalization for beta training frequencies and mu test frequencies (circa 8-12 Hz; see group statistical results for /ipa/ and /api/ faster rates. Although the mu/beta clusters do not achieve statistical significance for the slower speech rates, comparable clusters are evident in their group mean data. The individual results for S1119 are entirely comparable to the group mean data but show a much clearer distinction between the mu and beta bands (see especially plot for /api/ normal rate): These data show that mu training frequencies about 8-12 Hz generalise to beta band frequencies; and that beta training frequencies circa 20-30 Hz generalise to mu test frequencies. However, the plots also show a clear mu/beta discontinuity, with low classification accuracy for frequencies between roughly 13-20 Hz. This mu/beta discontinuity is not as evident in the group results, presumably due to individual differences in the precise frequency ranges of the mu rhythm (Pfurtscheller et al., 1997).

On a final note, we emphasise that we do not wish to imply from the foregoing that the supra-beta frequencies are simply “noise” frequencies. Gamma rhythms in the circa 60-80 Hz band are robust and functionally important features of motor-related MEG responses (e.g. Johnson et al., 2020; see also Cheyne, 2013; Cheyne & Ferrari, 2013); and high gamma band (> circa 80 Hz) rhythms are key dependent measures of brain function in electrocorticographic studies (Crone et al., 2006) including studies of speech-related brain function (e.g. Chartier et al., 2018; Conant et al., 2018). However in the context of a reiterated speech task any potential gamma frequency neural activity is strongly and visibly obscured by broadband movement artifact in both the time and frequency domains. From the foregoing analyses we conclude only that the mu/beta bands are suitable candidates for further interrogation of their functional properties, and that (within this experimental paradigm and MEG neuroimaging setup), the gamma band is not.

### Decoding of speech kinematic parameters from source- and frequency-constrained MEG data

## Methods

In the preceding analyses we have used standard MVPA classification of speech versus resting conditions to demonstrate that the neural signals derived from speech motor cortex contain information that is capable of discriminating between speaking and resting conditions; and then employed cross-frequency classification to determine that the mu-beta motor rhythms contain the informational basis of speech-rest discrimination. Importantly, cross-frequency generalisation also shows that the informational structure of the mu/beta rhythm is independent of the broadband artefacts that are an inevitable confound for electrophysiological recordings during overt speech.

In subsequent analyses we attempt to derive a more detailed picture of the information structures contained within the neural data, by performing classification between data partitions within the speech condition, rather than between speech and rest conditions. Representational Similarity Analysis (RSA; Kriegeskorte, 2008; Kriegeskorte & Kievit, 2013) is an MVPA technique based on the simple logic that classes of neural data with more similar informational (representational) structures should be more difficult to classify, relative to classes with more distinct representational structures. Previous studies have successfully applied RSA to tracking data of hand movements (Kolasinski et al., 2020), articulator movements during vowel production (Carey et al., 2017), and acoustic measurements during speech production (Zhang et al., 2020). We follow this logic to test specific hypotheses about potential representational structures in speech motor cortex activity as follows (see Figure 7 for a summary of the computational steps):

1. Our starting hypotheses concerning candidate representational structures within speech motor cortex activity come from the well-behaved kinematic profiles derived from direct MASK measurements of speech articulator movements (Figure 4): both the strikingly linear relationship between amplitude and velocity, and the orderly curvilinear relationship between duration and stiffness have been proposed to reflect “control parameters” that are relatively tightly specified at some level within the speech motor system (Munhall et al., 1989; Van Lieshout et al., 2007);
2. Within a given kinematic profile, we divide the behavioural data points into partitions (containing equal numbers of trials) that reflect different (Euclidean) distances between the coordinates within each partition. Here we have used 10 partitions which provides a good spread of inter-partition distances, each of which reflects a reasonable number of trials of MEG data (15-30 trials/partition).
3. Data points are averaged within each partition to provide a representation of the central tendency of each partition.
4. A “behavioural dissimilarity matrix” is generated based on the Euclidean distances between averaged data points in all possible pairs of partitions.
5. The MEG data (consisting of N trials by 3000-time points x 100 frequencies) is similarly divided into sets of individual trials that correspond to the behavioural data points within a partition.
6. Classification analysis is performed for all possible pairs of MEG trial partitions.
7. A “neural dissimilarity matrix” (for each time and frequency point) is generated based on the Euclidean distances between the classification accuracy scores.
8. An “RSA time-frequency plot” is generated containing the correlations between the behavioural dissimilarity matrix and the neural matrices for each time and frequency point.

Model evaluation is restricted in *time* to a 1 second epoch centred on the onset of the BC opening movement, as this event is the reference for the epoching of the MEG data. Model evaluation is further restricted to the *frequencies* of the mu and beta speech motor frequency bands of interest defined by the analyses described in the frequency localisation sections above; and for comparison purposes, a third high frequency region (60-80 Hz) dominated by speech movement noise and which we therefore do not expect to contain useable information concerning the representational structures of speech neuromotor activity.

Group data were statistically evaluated with cluster-based permutation analyses using 500 permutations, alpha < .05, and controlled for multiple comparisons using FDR.

The statistically significant results of model evaluation are shown in Figure 8. Of the two behavioural models, four speech conditions, and three frequency bands evaluated, only the /api/ normal rate condition shows showed statistically significant correlations, for beta band and for the stiffness-duration model. Three observations are relevant from these results: (1) The group mean correlations are overall very weak, with peaks restricted to a range of less than - .2 to .2; (2) The temporal structure of the significant positive clusters is appropriate, beginning at a latency of about 100 ms prior to the onset of the BC opening movement. This timing is in good accordance with what one would expect for neural activity associated with behavioural movements. (3) The post-movement cluster of correlations is oppositely (negatively) correlated to the premovement cluster.

Figure 8 also shows comparable (though non-significant after cluster-correction) results for /ipa/ normal rate for the velocity-amplitude model. While we do not wish to over-interpret a non-significant result, the circa -90 ms timing of the peak positive correlation cluster is entirely comparable to that observed for the /api/ normal rate stiffness-duration data described above.

Group mean behavioural-neural correlation time series are shown for all speech conditions, frequency bands, and behavioural models in Figure 9. As expected, in the gamma frequency band no significant results were obtained for any speech condition or behavioural model, and there is no discernible, consistent temporal structure in any of the plots. The mu band similarly shows no discernible or consistent temporal structure, and no significant results were obtained.

**Figure 9A, B, C.**
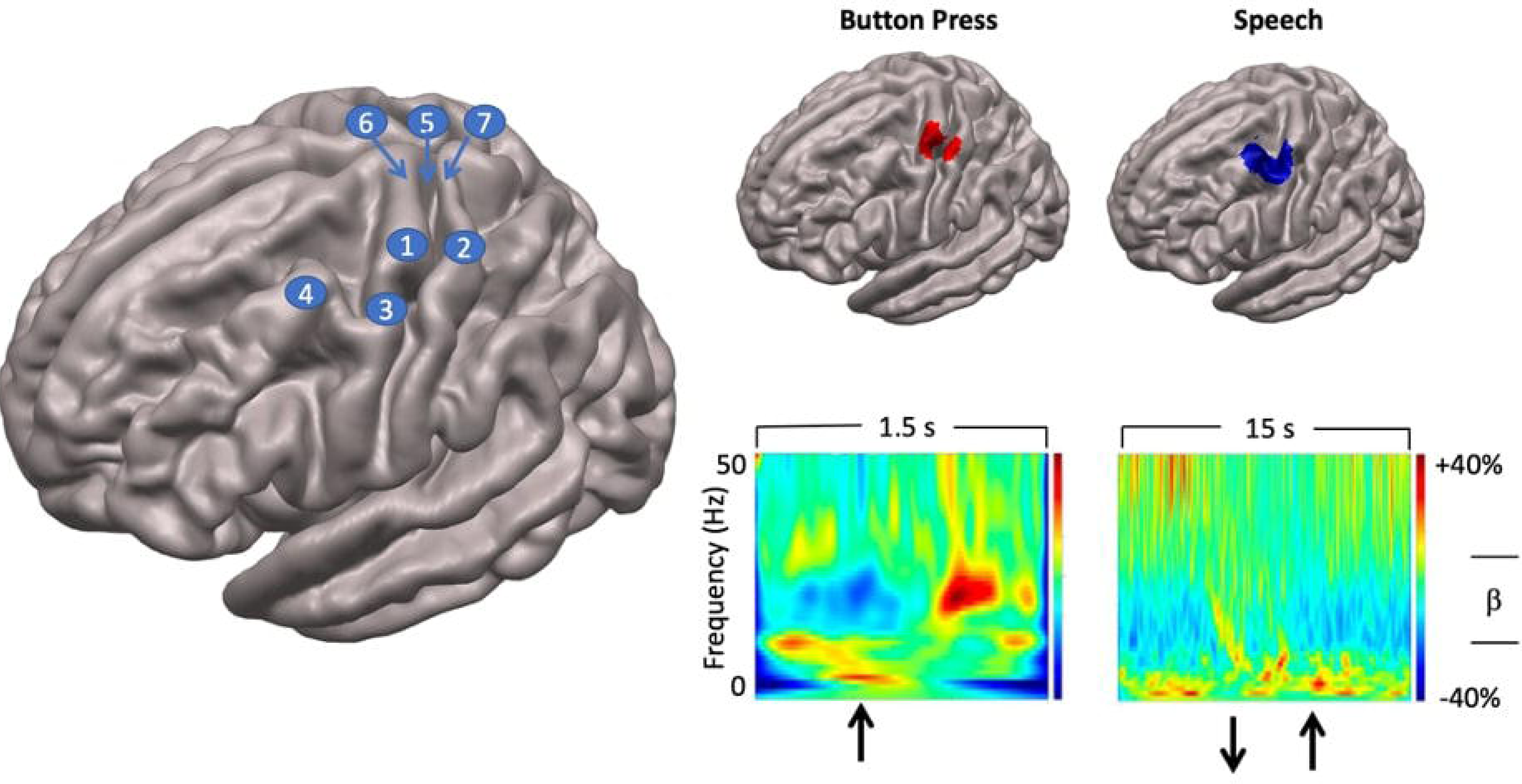
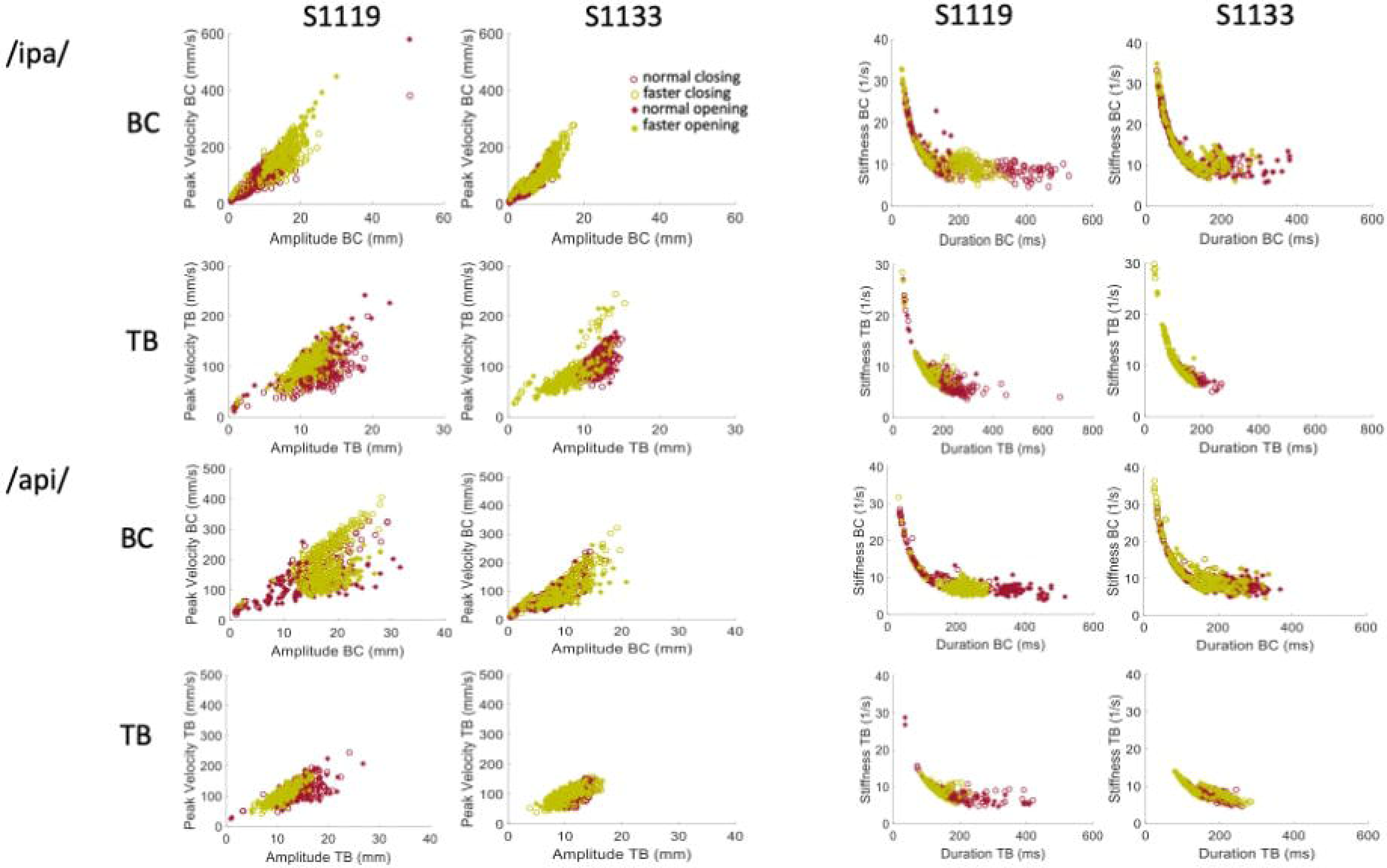
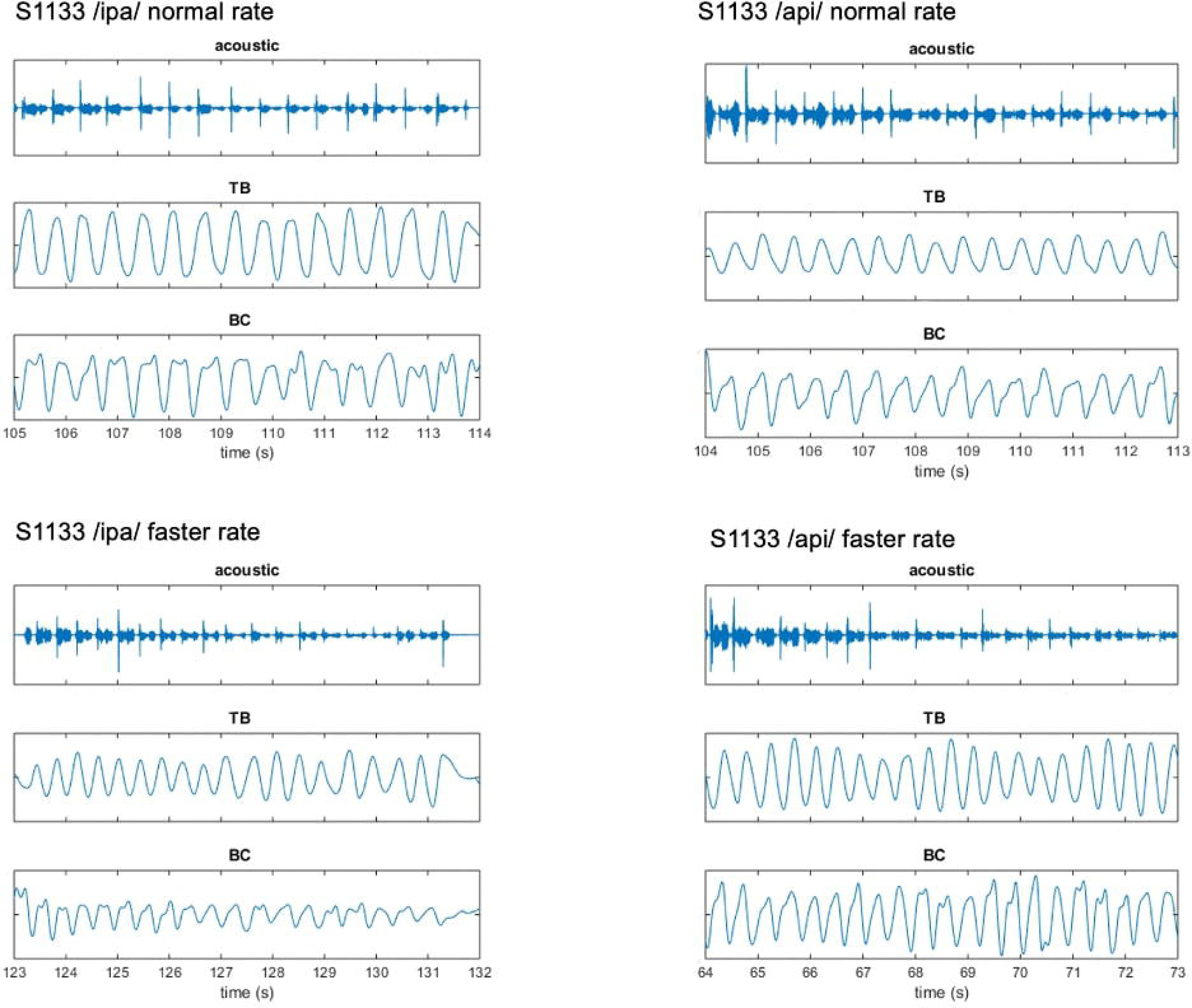
Group mean behavioural-neural correlations for all speech conditions, frequency bands, and behavioural models. Black lines show group mean correlations; shading shows standard errors.

In summary, the present results provide support for very weak but statistically significant encoding of the stiffness-duration relationship only in the beta motor rhythm. This encoding is statistically robust in only one speaking condition and is very weak in terms of magnitude of correlation. However, it is well-structured in time and shows an appropriate and expected temporal relationship (about -90 ms) with respect to movement onset.

## Discussion

### Localization of speech motor cortex

The results of our speech localiser analyses show a single statistically significant cluster in the left hemisphere, located immediately inferior to the hand motor cortex (hand knob) as identified with the independent button press localiser task. The cluster coordinates are centred on the posterior region of the middle frontal gyrus (pMFG) and extend posteriorly to encompass the middle portion of the precentral gyrus (midPrCG). This middle zone of the precentral gyrus – inferior to the precentral hand representation and superior to the ventral representations of face, mouth and larynx in the classical Penfield motor homunculus - corresponds to a functional-anatomical zone of the precentral gyrus that has been termed “dorsal laryngeal motor cortex” (dLMC; Bouchard et al., 2013; Eichart et al., 2020), a second representation of the larynx in addition to the ventral laryngeal motor cortex (vLMC) representation of the classical motor homunculus. The dorsal laryngeal representation appears to be unique to human motor cortex; this dual mode of organisation been proposed to underpin the extensive human capacity for voluntary control of vocalisation relative to non-human primates and other mammalian species (Belyk, Eichert & McGettigan, 2021).

The present results align with an emerging neuroscientific consensus which assigns novel and previously overlooked roles in speech motor planning and execution to the midPrCG and the immediately adjacent portion of the pMFG (area 55b; Glasser et al., 2016). Recent data from invasive neurosurgical and functional neuroimaging investigations have implicated the midPrCG in a variety of language and speech functions, ranging from direct laryngeal motor control to auditory-motor integration to higher level aspects of speech motor sequence planning that have previously been assigned to Broca’s area in models of speech processing (see reviews by Hickok et al., 2023, and Silva et al., 2022). Invasive neurostimulation of this area evokes involuntary vocalisations (Dichter et al., 2018). The midPrCG is anatomically contiguous with and tightly functionally associated with the pMFG, an area that has recently been implicated in the production of fluent speech (Glasser et al., 2016; Silva et al., 2022). Focal surgical resection of the pMGF and midPrCG has been reported to result in a case of pure apraxia of speech (Chang et al., 2020), a speech motor planning disorder associated with problems in hitting articulatory targets (articulatory groping) and transitioning between syllables. In light of these findings two recent theoretical models of speech production have posited a critical role for these areas in coordinating complex phonological sequences into motor plans, either in and of themselves (Silva et al., 2022), or as a subsystem of a dual (dorsal and ventral) motor speech coordination system (Hickok et al., 2023). Our results are clearly supportive of these models, since fluent reiterated (nonword) speech relies heavily on the capacity to coordinate and maintain the ordering of phonological sequences.

Invasive neurosurgical studies of midPrCG/pMFG function have focussed on the left hemisphere but there is currently little evidence to bear on the question of whether speech motor coordination functions of these areas have a lateralised or bilateral organisation (Hickok et al., 2023; Silva et al., 2022). The present speech localiser results showing a single significant cluster in the left hemisphere are generally consistent with those of a recent MEG study by DeNil et al. (2021), who (using the same analysis toolbox as the present study) compared beta-band responses to simple (a sequence of four repetitions of individual syllables /pa/, /ta/ or /ka/) and complex sequences (pseudo random combinations of four syllables, e.g. /patakapa/) of nonword vocal productions. These authors reported that beta-band event-related desynchronisation for both simple and complex verbal tasks was bilateral in premotor and motor cortical areas, but “statistically stronger” in the left hemisphere. We are aware of a single previous neuroimaging (fMRI) study using a continuously reiterated speech task that is overall comparable to ours. As in our study, Reicher et al. (2000) required participants to continuously produce nonword utterances during exhalation of a single breath intake. They used nonword items /ta/, /stra/ and /pataka/ that were consistent with German phonotactic rules but differed in articulatory/complexity. Consistent with our results, these authors reported highly focal activations of sensorimotor cortex for all task items. These activations were bilateral for the single syllable tasks, while the multisyllabic /pataka/ task showed activity only in the left hemisphere. Taken together, our results and those of Reicher et al. (2000) suggest that left lateralised speech motor function is associated with task requirements for more complex articulatory transitions in multisyllabic relative to monosyllabic utterances. A separate study by our group (Anastasopoulou et al., 2023 preprint) has further shown that typically developing children exhibit *bilateral* responses to the same disyllabic utterances used in the present study, indicating a developmental trajectory that shifts control of complex phonological sequencing from bilateral processing in childhood to left hemispheric dominance in adults.

### Decoding of speech-related brain activity

In recent years, a series of MEG studies have aimed to decode neural activity patterns associated with speech in the context of advancing the development of brain-computer interfaces (BCIs) (Dash et al., 2021; Dash et al., 2020; Dash, Wisler et al., 2020; Dash et al., 2019; Dash, Ferrari, Dutta, Wang, 2020). Using advanced artificial and convolutional neural network classifiers at the phrase level, Dash et al. (2020) have reported excellent classification accuracies for a five-class discrimination task (80% - >95%) for both imagined and overt speech, relative to comparable EEG studies achieving accuracies in the range of 35% (Cooney et al., 2019). A single previous MEG study (Dash et al., 2020) has also achieved accurate (average 80%) decoding of speech-associated jaw motion kinematics from concurrently acquired MEG signals, using a long short term memory (LSTM) model.

The present work makes three main contributions to this emerging body of MEG speech decoding literature. First, as discussed above, it provides a methodological and analytical approach for spatially constraining the overall decoding approach, effectively reducing the dimensionality of the problem from 160 MEG channels (or alternatively, thousands of source-reconstructed voxels) to a single virtual sensor. Further, there is now substantial evidence that the brain regions activated by our reiterated nonword speech task play a central role in in control and coordination of integrative speech movements, providing a highly plausible functional/anatomical candidate for more detailed interrogation of the kinematic representational structures that confer this control.

Second, the study incorporates a novel MASK technology which allows us, for the first time, to obtain detailed movement profiles of key articulators associated with the specific speech productions in our speech task in the same experimental setup as the MEG brain recordings and in temporal and spatial co-register with the brain data. Third, we describe an analytic pipeline using classification/decoding techniques that allow us to systematically query the nature the information contained in these kinematic representations, by testing specific hypotheses derived directly from overt movement behaviours. This analytic framework further permits strong inferences concerning the *timing* of relevant neural activations with respect to behavioural outputs.

Recent results from invasive ECoG studies of human neurosurgical patients provide compelling reasons to believe that concurrent neuroimaging speech tracking will be important for future progress in understanding speech motor control. For example, Chartier et al. (2018) obtained direct cortical recordings of human speech sensorimotor cortex together with (inferred) articulatory kinematics derived from a recurrent neural network based articulatory inversion technique which learned a mapping from produced speech acoustic to a speaker generic articulator space. This study showed that articulator movements were reflected significantly better in measured neural activity than were either acoustic or phonemic features of speech; that encoding is more related to coordinated movements of multiple articulators than to movements of single articulators; and that the behaviours of encoded movements were governed by damped oscillatory dynamics. These authors concluded that these coordinative and dynamical properties align neatly with the properties of articulatory units of speech (vocal tract gestures) as conceived within the theoretical framework of articulatory phonology and its associated task dynamics model (Browman & Goldstein, 1992; Goldstein & Fowler, 2003; Saltzman, 1986). As such, it seems clear that concurrent speech movement tracking and non-invasive neuroimaging should provide richer datasets with mutually reinforcing inferential power and precision relative to experiments that currently are largely conducted with only one or the other measure of speech motor control.

These new technical capabilities have clear clinical relevance for advancing our understanding and treatment of developmental and acquired disorders of speech. Speech-sound difficulties are the most common problems encountered by paediatricians and present formidable social, educational and employment obstacles in cases where these problems cannot be readily treated and resolved (Morgan, 2018). Childhood apraxia of speech (CAS) is an intriguing example of a highly debilitating and persistent disorder of speech development whose origins are considered to lie within the brain mechanisms responsible for coordinating and sequencing speech movements, but whose study with conventional neuroimaging approaches has so far proved highly resistant to establishing any clear connection to any particular brain region. In such cases, the capability to directly map speech kinematic and coordination function in speech motor control centres within highly focal and specific brain regions promises to provide more powerful insights into the origins of speech problems in CAS (and conversely, into why speech development proceeds more smoothly in most other children). Similarly, acquired apraxias of speech are a common and debilitating outcome of strokes and other brain injuries. The greater functional specificity of MASK-MEG has a clear bearing on studies aimed at understanding the nature and degree of functional compromise and plastic capabilities in the brain of these patients.

### Limitations

While our analytic framework dramatically and effectively reduces the dimensionality of the MEG analytic problem, there remains a large decision space concerning the selection of speech behaviours for input to the overall analyses. Here we have fairly arbitrarily focussed on a single articulator metric (the first opening movement of the bilabial closure gesture) and have tested 2 simple models (derived from the observed kinematic profiles) of the neural informational structures that may underlie this movement. Our results nonetheless provide support for the conclusion that the stiffness-duration relationship of the first opening movement of the bilabial closure may be (very weakly) encoded in the beta-band sensorimotor rhythm, with a timing beginning approximately 90 ms before the onset of the movement. The reason for the occurrence of a second period of significant (but oppositely valanced) association with stiffness-duration at a latency of about 200 ms post movement is presently unclear: one possibility is that reflects a sensory reafference event that provides a check on the motor commands.

Overall, since we have obtained only a weak association between speech behaviour and neural activity (and in only one of four speech conditions), it is clear that future work should more systematically probe the sets of possible models of speech movement encoding, including models that describe relationships between articulators that are likely required for integrative speech behaviours (e.g., the Linguistic Gestural Model (LGM) which is a combination of Articulatory Phonology and Task Dynamics (Saltzman & Munhall, 1989; Browman & Goldstein, 1992; Browman & Goldstein, 1997, the speed-accuracy trade off known as Fitts’ law (Fitts, 1954; Neufeld & Van Lieshout, 2014; Gafos & van Lieshout, 2021; Kuberski & Gafos, 2021).

Finally, the present decoding framework employs a simple binary discriminant classifier and a simple RSA correlational mapping approach, which we have chosen for their ease of use and the straightforward inferential capabilities. At the present time artificial and convolutional neural network classifiers are undergoing rapid development with dramatic improvements in accuracy (see Dash et al., 2021; Dash et al., 2020; Dash, Wisler et al., 2020; Dash et al., 2019; Dash, Ferrari, Dutta, Wang, 2020 for recent examples with application to MEG speech decoding) and we anticipate that these improvements will be able to accordingly extend the accuracy and scope of MASK-based studies of speech motor control.

### Conclusions

MASK-MEG addresses an important gap in current neuroscientific capabilities for studying expressive language function in the human brain. While we possess robust and well-established methods for measuring and characterising overt movements of the speech articulators, and highly sophisticated equipment and methods for defining the brain activities that control these movements, the two methodologies are not readily or easily combined within a single experimental setup. As a result, speech movement tracking and speech neuroimaging methods have largely evolved within separate laboratories --even separate disciplines -- and there remains no easy way to co-register and reconcile the different types of information that are derived from them. The advent of neuroimaging-compatible speech tracking technologies such as MASK opens a new window for integrative studies of human speech motor control, combining precision measures of overt speech behaviours with temporally co-registered and spatially localised measures of brain function and new machine-learning based decoding approaches capable of interrogating the kinematic information structures represented in speech motor cortex.

## Author Contributions

IA conducted the experiment. IA and BJ analyzed the data. All authors conceived, designed the experiment, discussed the results, wrote, and edited the manuscript.

## Data Availability Statement

The raw data supporting the conclusions of this article will be made available by the authors, without undue reservation.

## Supplementary Materials

**Figure S1.**
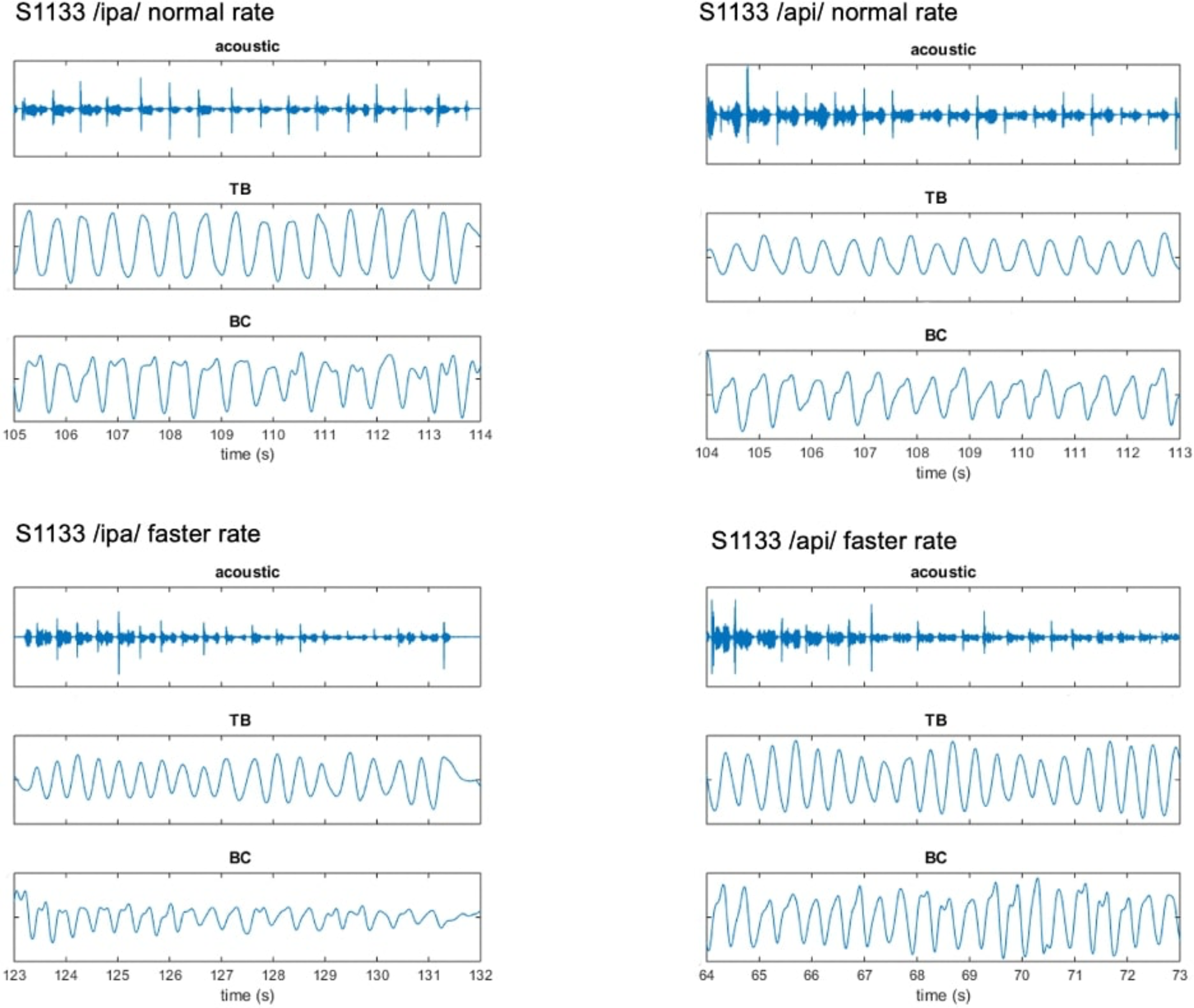
Representative acoustic and kinematic measurements from MASK. Data are shown for two participants for a single /ipa/ trial set at normal and faster speaking rates. Shown are (from top to bottom) waveforms for the audio signal, tongue body (TB) gesture, and bilabial constriction (BC) gesture.

**Figure S2.**
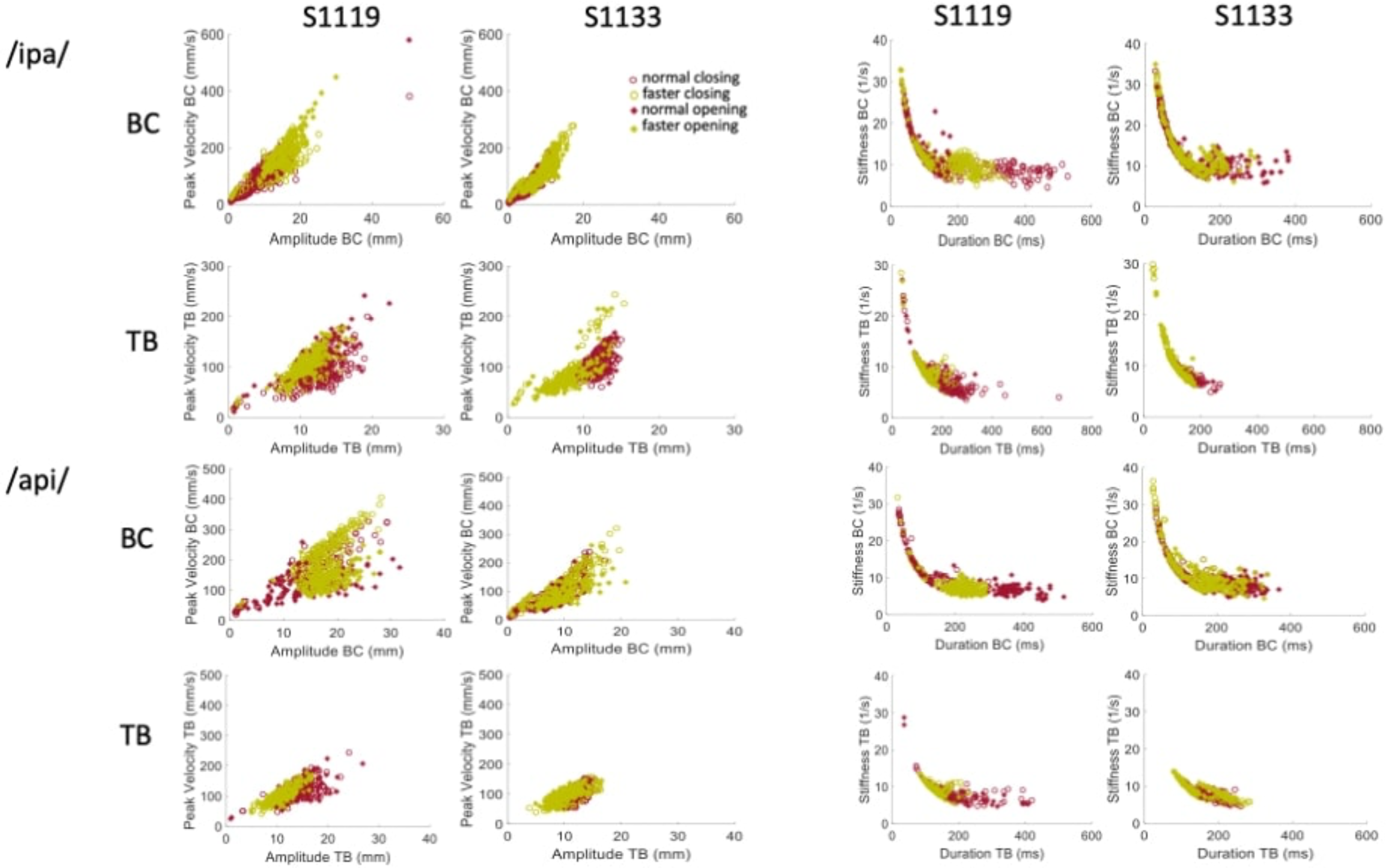
Covariation of kinematic parameters of speech movements for two participants. Left columns: Velocity versus amplitude. Right columns: Stiffness versus duration. BC = bilabial closure. TB = tongue body.

## Footnotes

1 In the case of event-related experimental designs the time-frequency classification can also determine if discrimination is confined to specific times (Treder, 2020). The reiterated speech paradigm used here is akin to a system in steady state, so the analytic question at this stage simplifies to frequency discrimination alone.

